# Characterizing MINFLUX imaging performance with DNA origami

**DOI:** 10.64898/2026.02.24.707670

**Authors:** Alexander H. Clowsley, Alexandre F. E. Bokhobza, Radoslav Janicek, Karol Kołątaj, Gabriela Bleuer, Lorenzo Di Michele, Guillermo P. Acuna, Christian Soeller

**Author notes:** Corresponding authors &.

## Abstract

MINFLUX, a second-generation super-resolution technique, can localize fluorescent markers approaching single-nanometer precision in three dimensions. Similar to previous super-resolution methodologies, the extended duration of acquisitions can result in drift that needs accurate correction to enable proper analysis and interpretation of the collected data. Here, we use DNA origami, housing sites of known spatial distribution fitted with repeat-domain docking strands for DNA-PAINT imaging to characterize imaging performance over extended duration MINFLUX acquisitions (6-20 h). Repeat-domain docking strands overcome site-loss and reveal residual drift in prolonged MINFLUX 3D acquisitions that we correct with an algorithm exploiting time-correlated shifts of localizations around identified DNA origami sites. Following correction of residual drift the site precision, i.e. the scatter of localizations around sites, is ∼2 nm in all directions. Comparison of site precision from extended repeat-domain docking strands with site precision from standard short 8-10 nucleotide docking strands exhibits no detectable loss of site precision. By adding DNA origami structures to mounted biological samples we apply our approach to the imaging of the cardiac ryanodine receptor 2 in cryosectioned heart tissue. The data suggests that for these protein targets single domain markers with repeat domain docking strands may be directly used for residual drift correction, simplifying sample preparation and acquisition protocols.

## Introduction

Recent super-resolution approaches pushing the resolution limit towards the nanoscale have largely revolved around two ideas: engineering the point-spread function (PSF), as with STED,^1,2^ or by isolating single emitters by stochastic mechanisms, as with STORM/PALM^3,4^ and related SMLM (single molecule localization microscopy) methods. MINimal fluorescence photon FLUXes (MINFLUX) ‘nanoscopy’ combines these two methodologies to achieve isotropic resolution in 3D^5^, where a principal advantage stems from using a minimum in the PSF to localize emitters^6^. For 3D imaging, a bottle-beam with a zero-intensity center is rapidly scanned across regions of interest scouting for single-molecule events. These single-molecule events are typically achieved through stochastic switching of photo-switchable fluorescent probes, like Alexa Fluor 647, or by the transient interaction of complementary oligonucleotide pairs in an approach called DNA-PAINT.^7^ In DNA-PAINT, the location of interest is pre-labeled with a short docking site where a complementary dye modified ‘imager’ oligo can transiently interact. This method helps to alleviate the requirement to use special chemical switching buffers. DNA-PAINT has proved versatile in solving various imaging challenges such as those from non-specific interactions,^8,9^ high backgrounds,^10^ and has additionally enabled the creation of a number of super-resolution nanotools for measuring cellular traction forces,^11^ and nanometer proximity sensors.^12^ Despite circumventing the effects of photo-bleaching, which are prominent in photo-switching experiments, DNA-PAINT does suffer from docking site-loss^13^, an effect that may be exacerbated by the serial detection of current MINFLUX implementations.

Acquiring super-resolution images is time consuming and with the serial detection of current MINFLUX implementations it can take several hours to obtain a single dataset. This is especially true if care is taken to harvest the information in densely labelled samples as we have recently systematically investigated^14^. Inevitably this makes drift, and its correction, a critical factor in MINFLUX experiments, which needs to be considered to fully exploit the obtainable localization precision. We therefore sought to establish a method to characterize drift in extended duration (>1h) MINFLUX acquisitions that to our knowledge had not been carried out before. To this end, we adapt the use of super-resolution templates made from DNA origami^15,16^ to characterize MINFLUX imaging performance. To function as super-resolution imaging templates DNA origami are designed to include fluorochromes^17^ or, for DNA-PAINT, docking sites for fluorescent imager strands at known locations and spacings^7,18^. Critical for the characterization of long-term MINFLUX imaging performance demonstrated here is the introduction of repeated docking domains at DNA origami sites. We and others had previously introduced repeat DNA-PAINT^19^ for SMLM. For both test DNA origami samples and biological samples^14^, Repeat DNA-PAINT with extended docking domains allows overcoming DNA-PAINT site loss and is compatible with many repeat visits at single anchor sites. We introduce the use of these DNA origami structures, harboring repeat docking domains, as calibration tools for MINFLUX acquisitions to characterize and correct residual drift that remains after employing built-in stabilization and drift removal systems, and illustrate this with a commercial MINFLUX system in our laboratory. We also confirm experimentally that there is no detectable spatial broadening from the use of “long” (>40 nt) repeat-domain sequences on DNA origami compared to conventional “short” (8-10 nt) docking domains. Finally, we demonstrate that by incorporating repeat-domain DNA origami with biological samples it becomes possible to quality assure long-term MINFLUX 3D imaging and get closer to the limiting performance set by MINFLUX localization precision.

## Results

To demonstrate the suitability of DNA origami with repeat-domain docking sites for long-term 3D MINFLUX imaging, two DNA origami template types containing 6 repeat-domain docking sites per structure (10xRD P1 docking design, see Table 1) were imaged for several hours. Figure 1a-i shows a rendered MINFLUX overview image that shows many DNA origami tiles with the 6 anchor sites well-resolved. In a close-up of 3 nearby tiles in in-plane and side-on y-z views (Fig. 1a-ii and 1a-iii), repeated visits to anchoring sites are apparent (each dot is obtained from a single MINFLUX “trace” that was recorded, see also methods) with some spatial scatter around each site discernible. The DNA origami tiles (Design O1) used in this sample have anchor sites on a ∼45 by ∼40 or 35 nm grid spacing as shown in the schematic in Fig. 1a-iv. The statistics of site visits confirm that anchor sites are visited repeatedly (on average 14.5 times, Fig. 1a-v) and the rate of MINFLUX acquisitions stays approximately constant at ∼150 traces per 400 s throughout an ∼11 hour acquisition, indicating that site-loss can be effectively overcome with the repeat-domain approach. We conducted similar experiments using a second DNA origami design O2, also containing 6 anchor sites (Fig. 1b-i with details views of repeated visits at anchor sites in Fig 1b-ii and b-iii) but on a tighter 15 by 20 nm grid spacing. Use of repeat-domain docking strands at the anchor sites similarly ensured many repeat visits (here on average 10.2 times, Fig. 1b-v) with a near-constant localization rate of ∼150 traces per 400 s throughout an ∼8.5 h acquisition.

**Figure 1.**
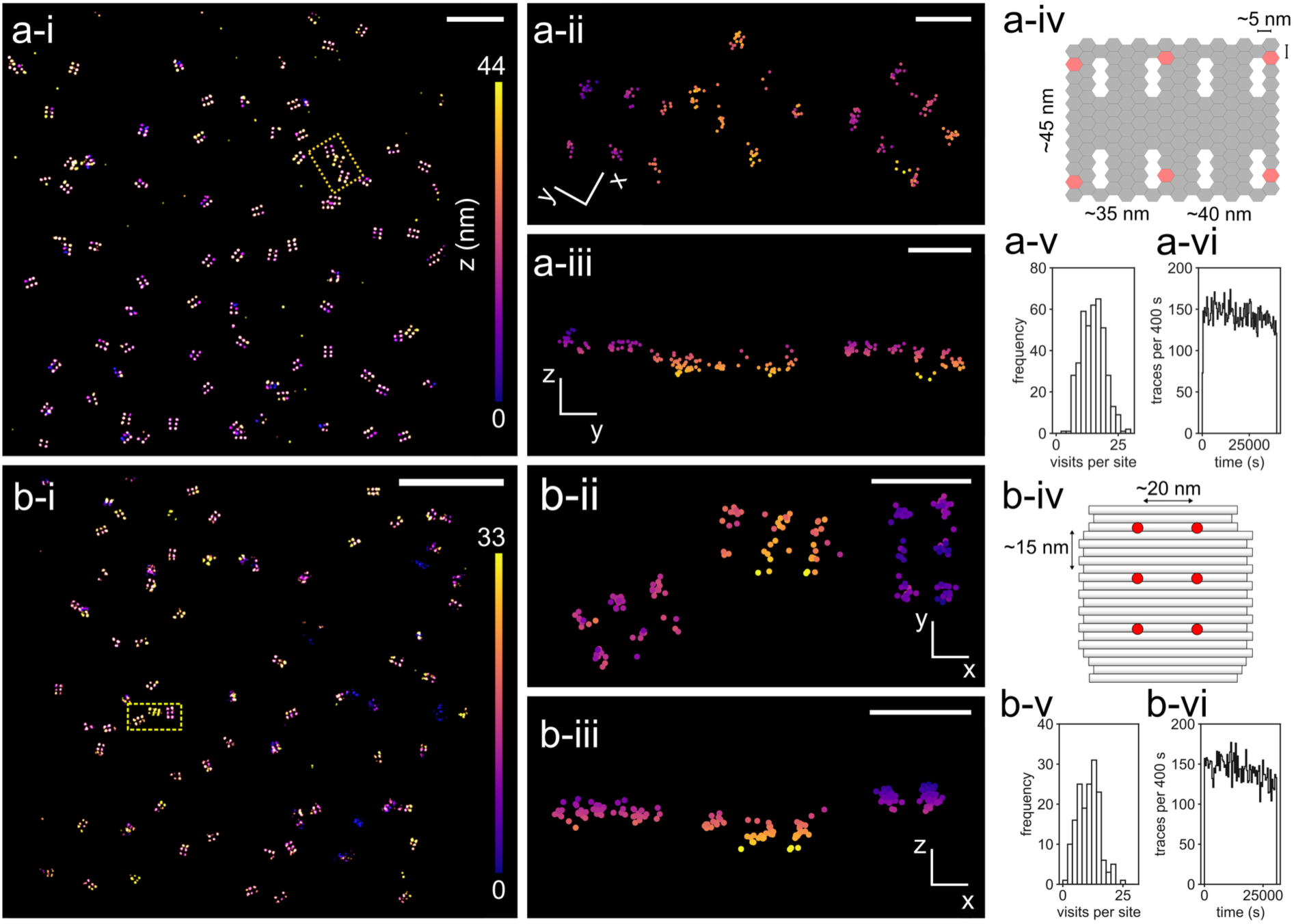
Long-term 3D MINFLUX imaging of DNA origami. **a-i**.MINFLUX overview image showing DNA origami structures O1 functionalized with repeat domain docking strands. **a-ii, iii**. Detail views of the yellow framed area in a-i in the x-y and y-z planes, respectively. Each dot shows a coalesced localization with color according to z-coordinate. **a-iv**. Template design O1 showing placement of anchor strands. **a-v**. Histogram of site visits during the 4·10^4^ s duration (11.1 h) acquisition with 14.5 visits on average. **a-vi**. Rate of localization quantified as number traces per 400 s. b-i. MINFLUX overview image showing DNA origami structures O2 functionalized with repeat domain docking strands. **b-ii, iii**. Detail views of the yellow framed area in b-i in the x-y and y-z planes, respectively. Each dot shows a coalesced localization with color according to z-coordinate as in b-i. **b-iv**. Template design O2 showing placement of anchor strands on 15×20 nm grid. b-v. Histogram of site visits during the 3.1·10^4^ s duration (8.6 h) acquisition with 10.2 visits on average. **b-vi**. Rate of localization quantified as number traces per 400 s. Note the stable localization rate over the full duration of acquisition. The localizations shown were corrected by the MBM system using all MBM beads that were monitored (see also methods). Scale bars a-i, b-i: 500 nm; a-ii,a-iii,b-ii,b-iii: 50 nm.

**Table 1.**
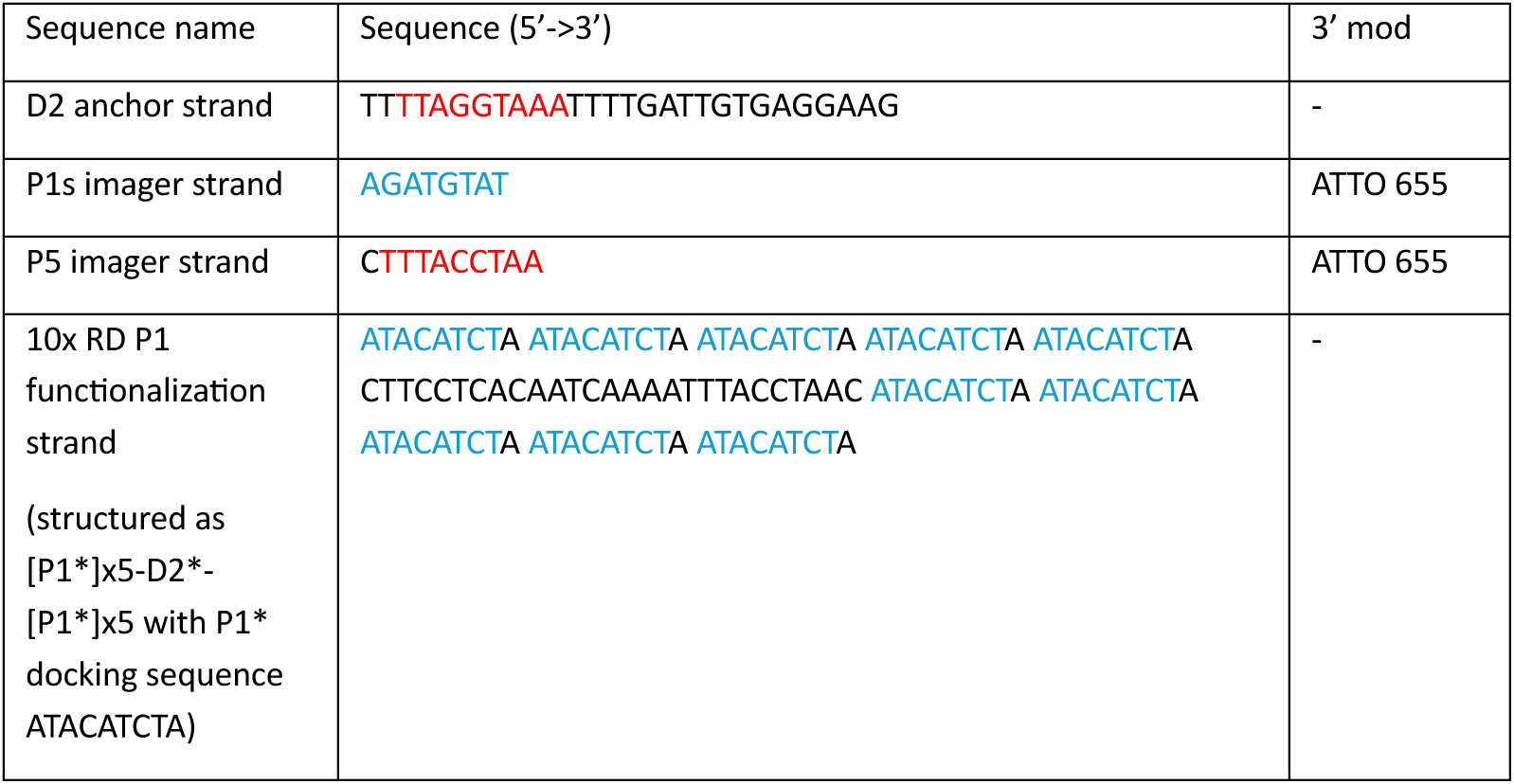
DNA-PAINT sequences used. The complementary P5 imager and P5* docking domains are colored red. The complementary P1s imager and P1* docking domains are colored blue.

We used DNA origami MINFLUX data as in Fig. 1 to characterize the imaging performance of the commercial MINFLUX microscope. Figure 2a shows the gradual drift present in a MINFLUX dataset where localizations were not corrected for MINFLUX beam alignment. In our experience, MINFLUX beam alignment is the major remaining source of drift in scanning MINFLUX microscopy when the sample is actively stabilized on the microscope stage to better than 1 nm, as in the commercial system characterized here. The resulting drift can be gleaned from the systematic displacement of site localizations that is apparent when these localizations are colored by acquisition time, see Fig. 2a. The majority of this drift is eliminated when MINFLUX beamline monitoring (MBM) information is used to correct the localization positions (Fig. 2b). The MBM system tracks gold nanoparticle fiducials throughout the acquisition period, approximately once every second, and from this information estimates drift trajectories in x, y and z-directions (for 3D MINFLUX acquisitions) that are shown in Fig. 2c. The distribution and selection of nanoparticles for correction, the nanoparticle localization processing and generation of MBM drift trajectories are shown in Supplementary Fig. 1 and described in the supplementary methods. While the MBM correction greatly reduces the visible drift we sought to estimate any remaining residual drift from the knowledge that spatially tightly grouped localizations originate from single anchor sites on DNA origami. We implemented an algorithm that estimates residual drift from the time-correlated dispersion of localizations around anchor sites (we term the approach “correlated site-dispersion”) as described in the methods section on data analysis and illustrated in Supplementary Fig. 2.

**Figure 2.**
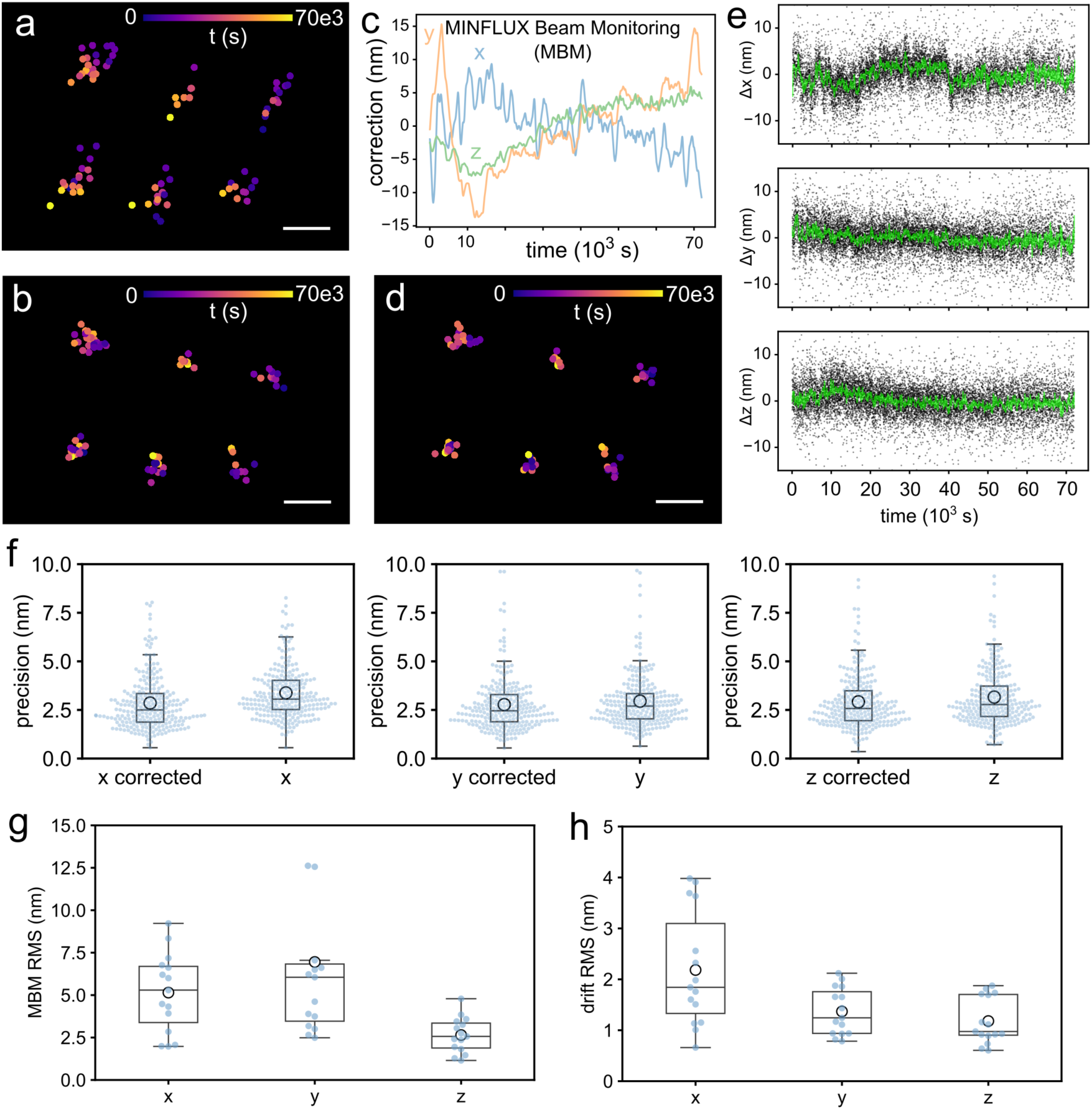
Determination and reduction of residual drift with DNA origami. **a**. MINFLUX localization from a single DNA origami tile without any drift correction. Localizations are shown as coalesced localizations (see methods) colored by time of recording. **b**. The same localizations after drift correction using the mean MINFLUX beamline monitoring (MBM) trajectories, where the MBM trajectories are shown in panel **c**. **d**. After additional residual drift correction based on correlated site-dispersion (see methods and estimated trajectories in panel **e**), the localizations appear visually tighter packed around the site centroids as compared to MBM-only correction (shown in b). **f**. Quantitative analysis of the site precision (measured as the standard deviation of site group coordinates) using the dataset from which the localizations in a-d were extracted, demonstrates an improvement in median precision by ∼0.2 nm in y and z directions while the improvement in x was ∼0.6 nm (249 sites from a single MINFLUX 3D dataset). **g**. Root mean square (RMS) amplitudes of the MBM measured drift were between ∼2.6 (z) and 6.1 nm (y) on average (n=15 data sets from N=6 independent replicates). **h**. Corresponding RMS amplitudes of residual drift were observed to be typically in the range of 1-2 nm on average (1.8, 1.2 and 1.0 nm in x, y and z directions), with largest values approaching 4 nm in the x-direction, n=15 data sets from N=6 independent replicates. Scale bars a, b, d 20 nm.

Subtracting the estimated residual drift from the localization positions resulted in a tighter distribution of localizations around the DNA origami sites, as shown in Fig. 2d. The corresponding residual drift trajectory estimates in x, y and z directions are shown in Fig. 2e. In this dataset, the largest drift amplitudes are seen in the x-direction, with residual drift amplitudes up to ± 5 nm over a >70k s acquisition (∼20 h). Residual drift in y and z-directions is comparatively smaller but non-negligible given the high localization precision of MINFLUX acquisitions. To measure the improvement resulting from residual drift correction, we quantified the site-precision determined as the standard deviation in x, y and z directions of groups of localizations originating from single origami sites. Groups of localizations from single anchor sites were identified by DBSCAN clustering^20,21^ with a neighborhood radius ε = 15 nm given the well-spaced sites of the origami design with sites at least ∼35 nm apart (see also Fig. 1a-iv). For the MINFLUX dataset shown in Fig. 2a-d, the correction using the estimated residual drift trajectories improved site precision from 3.1 nm (MBM correction-only) to 2.5 nm (median) in x, from 2.7 to 2.5 nm in y, and from 2.8 to 2.6 nm in the z direction (Fig. 2f), i.e. the corrected data has near isotropic precision of ∼2.5 nm in all directions. We carried out a range of similar experiments and observed that the drift corrected by the MBM system had root mean square (RMS) magnitudes of 2.6 (median in z) to 6.1 nm (median in y) over a period of up to 30k s (∼8 h), as shown in Fig. 2g (n=15 data sets from N=6 independent replicates), while the residual drift RMS magnitudes were between a median of 1.8 (x) and 1.0 nm (z, Fig. 2h, n=15 data sets from N=6 independent replicates). When quantified as the total range, i.e. the difference between most positive and most negative drift distance, the residual drift total range was between ∼10 nm (median in x) and ∼6 nm (median in z), where the residual drift range in the x-direction had generally the greatest magnitude, see Supplementary Fig. 3a (n=15 data sets from N=6 independent replicates), with associated MBM recorded total drift ranges between ∼28 and ∼12 nm (Supplementary Fig. 3b, n=15 data sets from N=6 independent replicates). Supplementary Fig. 3c-d shows an example MINFLUX acquisition with a more marked initial residual drift over the first ∼12000 s. Correction of this residual drift improved the Fourier ring correlation (FRC), a measure of the lateral resolution, from 11.6 to 6.2 nm (Supplementary Fig. 3g, h) once the residual drift was corrected and data from only those initial 12000 s were used for rendering. Similarly, the Fourier shell correlation (FSC), a measure of the 3D resolution, was 6.3 nm (Supplementary Fig. 3i) indicating that the residual drift correction also works efficiently in the z-direction. Generally, larger drift, as measured by the MBM system, was correlated with more pronounced temperature changes as illustrated by the corresponding signal from a temperature sensor at the microscope stand (Supplementary Fig. 3j). Overall, the data shows that DNA origami with repeat-domain sites allows estimating residual drift and that correction of this residual drift (additional to the MBM system based correction) improves site precision and spatial resolution measures.

A key to enabling long-term recordings from individual DNA origami anchoring sites with 3D MINFLUX is the use of extended repeat-domains, here using an extended strand containing 5 P1 domains on either side of a central tethering domain (see Table 1). Accordingly, each single stranded (ss) domain extends over 5×9, i.e. 45 nt, which would correspond to 45 x 0.33 nm, or 14.9 nm, when stretched out in a double-stranded configuration. We had argued previously that the rapid thermal fluctuations of the mostly ss DNA occurring on a time-scale of microseconds of the extended repeat-domain (Fig. 3a), should essentially average out over the recording time (typically several ms in MINFLUX localizations) and lead to negligible loss of localization precision^19^. We tested this idea by comparing the site precision (see repeatedly visited sites in Fig. 3b) from unmodified anchor sites exposing docking site D2 (see Table 1), containing a short 9 nt P5* domain (Table 1), with the site precision from MINFLUX recordings where 10x RD P1 repeat-domains (see Table 1) had been attached. Prior to this comparison, residual drift was corrected for each dataset as shown in Fig. 2. The histograms of site precisions from unmodified sites (Fig. 3c, n=13 MINFLUX acquisitions from N=2 independent replicates) and repeat-domain sites (n=9 MINFLUX acquisitions from N=2 independent replicates) look comparable. When compared statistically, the differences in site precision across all coordinate axes were non-significant between unmodified and repeat-domain sites (p>=0.05, two-tailed T-test for the means of independent samples, n=8 acquisition series (10xRD) versus n=13 series (no modification), from N=2 technical repeats each) with mean values approximately 2 nm in all directions (Fig. 3e). Notably, site-loss with the unmodified single docking domain (P5) and Atto 655 conjugated imagers was substantial so that typically few complete origami tiles were recorded (Fig. 3f) whereas MINFLUX imaging with repeat docking domains (10x RD P1 and Atto 655 conjugated imagers) was compatible with extended duration MINFLUX imaging and acquisition of many complete DNA origami tiles (Fig. 3g). This difference is also apparent when the steep decay of localization rate with single docking domains is compared with the almost undiminished localization rate after > 8h of 3D MINFLUX imaging with 10x RD P1 repeat-domains (Supplementary Fig. 4), similar to prior observations with biological samples^14^.

**Figure 3.**
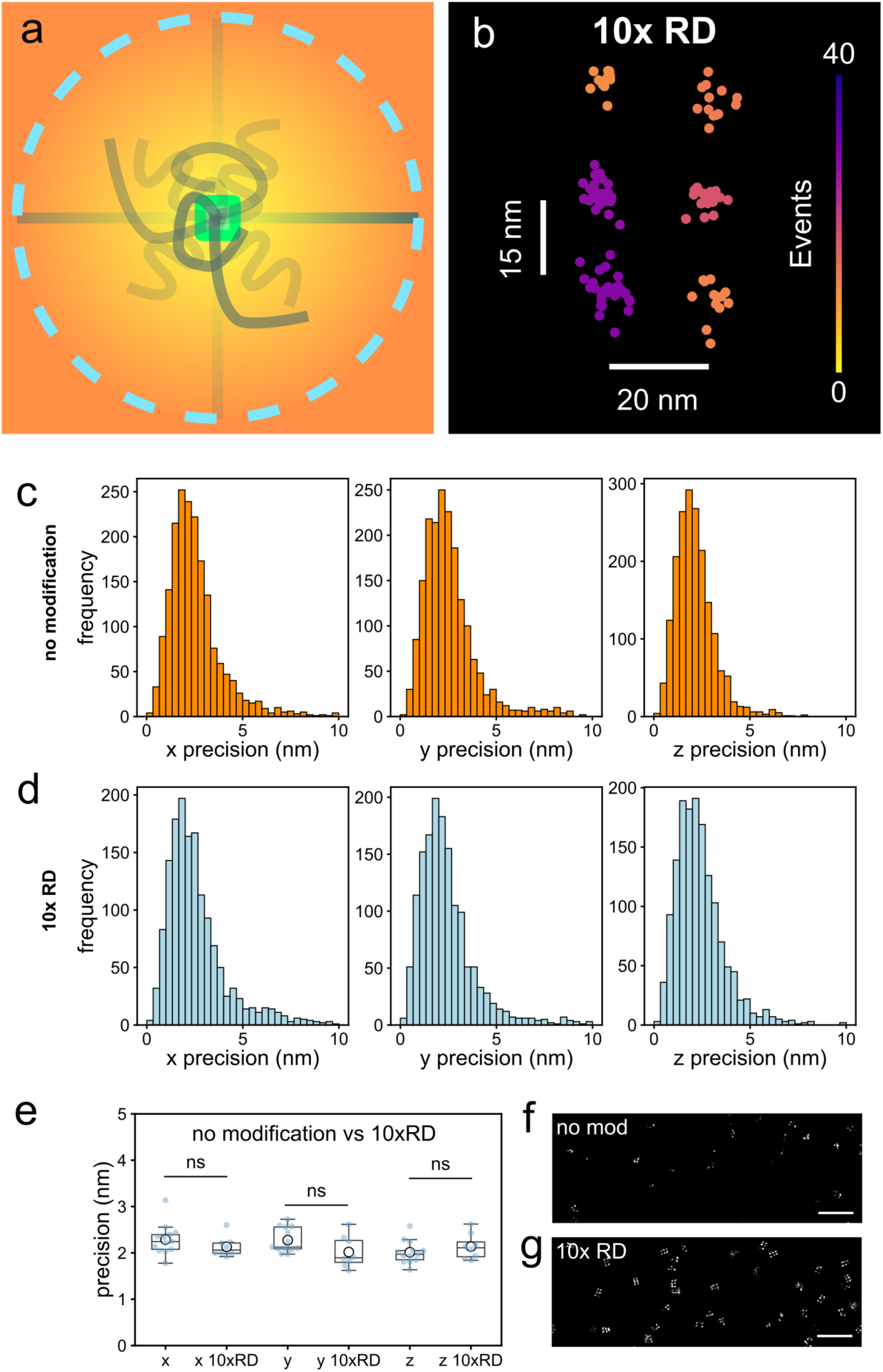
Localization precision of extended repeat-domain sites is indistinguishable from standard docking domain precision. **a**. The single-stranded docking domains of the repeat domain strand are expected to undergo rapid random coiling. **b**. Example of the distribution of coalesced MINFLUX localizations around the 6 repeat domain sites on DNA origami structure O2. **c**. Histograms of site precision from docking sites probed with no modification with imager P5 (n=13 MINFLUX acquisitions from N=2 independent replicates). **d**. Equivalent histograms of site precision from docking sites with extended repeat domain strand 10x RD P1 probed with imager P1 (n=9 MINFLUX acquisitions from N=2 independent replicates). **e**. Statistical comparison between MINFLUX data from unmodified anchor sites versus 10xRD repeat domain sites exhibits no significant difference in all axes (p>0.05, n=13 MINFLUX acquisitions (no modification) versus n=8 acquisitions (10xRD) from N=2 independent replicates). **f**. Site-loss is pronounced when rendering image data acquired with unmodified anchor sites whereas (**g**) with 10x RD repeat domains the DNA origami patterns are much more complete showing that repeat domains enable long-term MINFLUX acquisitions. Scale bars f, g 250 nm.

Having established the use of repeat-domain DNA origami for long-term MINFLUX imaging and correction of residual drift (with negligible penalty from the use of extended repeat docking domains) we sought to include these DNA origami as reference structures with 3D MINFLUX imaging of biological samples. This would enable “quality-assured” MINFLUX acquisitions of biological samples and allow residual drift removal. As a suitable biological sample, thin (3-5 µm thickness) cardiac tissue sections of ventricular tissue were used. These were taken from the hearts of PA-RFP RyR2 mice in which RyR2 (ryanodine receptor type 2) subunits are tagged with PA-TagRFP^22^. To label these for MINFLUX microscopy, they were stained with DNA-PAINT sdABs (single-domain antibodies) against tagFP^19^, and repeat-domains (10x RD P1) were attached, similarly to what was previously done for MINFLUX imaging of proteins in the nuclear pore complex^14^. Finally, DNA origami were added with their anchor sites also attached to the same 10x RD P1 repeat-domains. Imaging was conducted with P1 Atto 655 imagers which could reversibly hybridize both with the DNA origami repeat-domain sites and with the sdAB attached repeat-domain docking strands on RyR2s. When MINFLUX acquisition ROIs were placed on ventricular cells (guided by fluorescence from anti-tagFP sdABs conjugated to Alexa 488, Fig. 4a) the resulting MINFLUX images contain both MINFLUX localizations from RyR2s and DNA origami, which can be distinguished based on their morphology, Fig. 4b. During the 43k s acquisition (∼12 h) of the MINFLUX dataset shown, the MBM system tracked drift of up to 60 nm (Fig. 4c). The MBM corrected data was subjected to the residual drift estimation approach using the DNA origami data as illustrated above in Fig. 2 and Supplementary Fig. 2. This improved the tightness of localizations at origami sites which is apparent by comparing Fig. 4d and 4e, with the corresponding residual drift trajectories shown in Fig. 4f. Interestingly, some of the “planar” DNA origami templates adhered to the biological sample at various angles, such that some templates were oriented approximately vertically as illustrated with the DNA origami tile in Fig. 4g. These vertically oriented DNA origami directly confirm both well-corrected drift and high spatial resolution in the axial direction.

**Figure 4.**
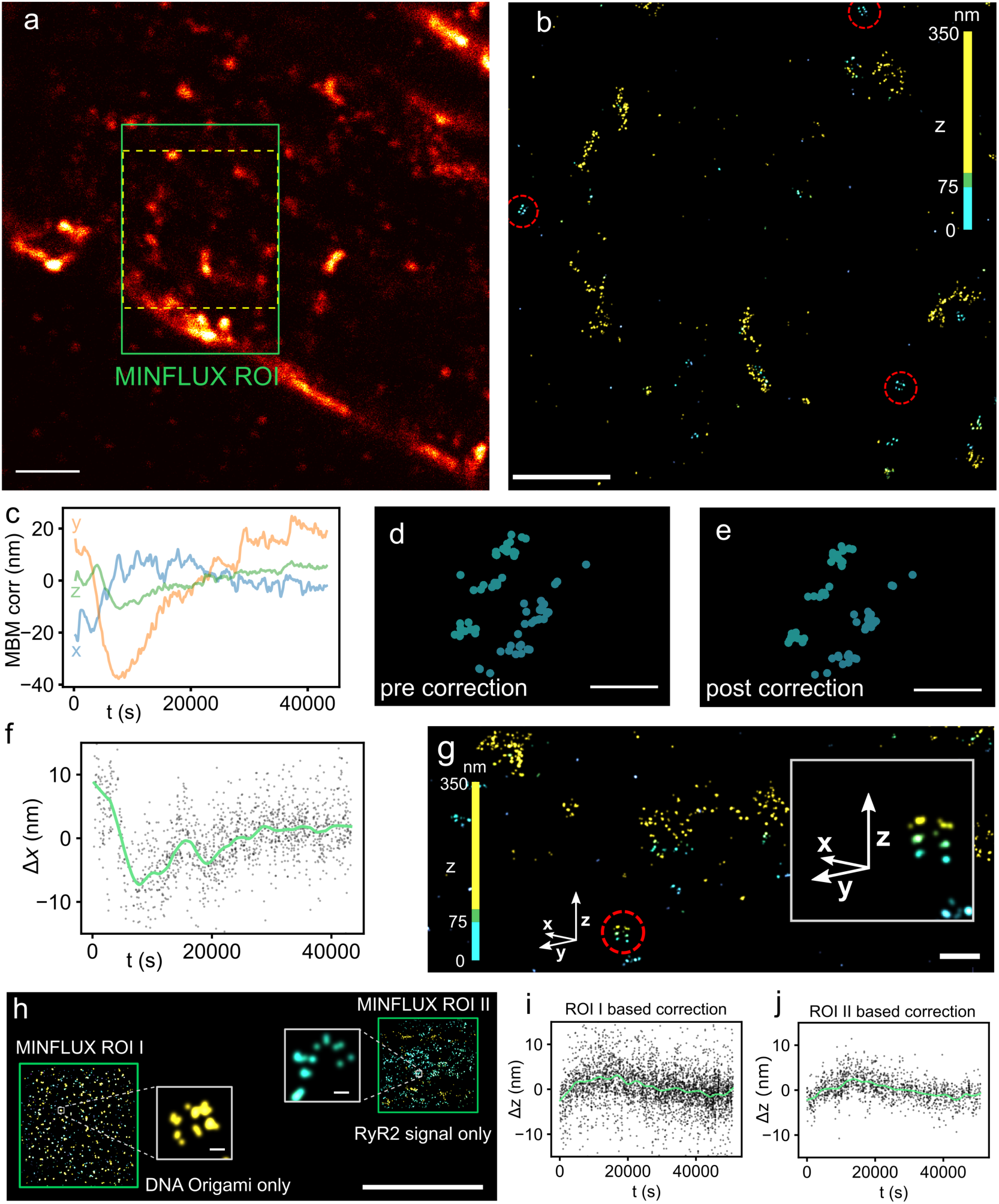
MINFLUX imaging of DNA origami with RyR2 labeling in cardiac tissue slices for quantifying residual drift. **a**. Confocal image of a tissue slice from a heart containing PAtagRFP-RyR2 counterstained with anti-tagFP Alexa 488. DNA Origami structures O1 were added to this sample (see methods). The green rectangle shows the ROI for MINFLUX 3D imaging. **b**. MINFLUX dataset rendered with PYMEVisualize and colored for z coordinate (region shown corresponds to the yellow dashed outline in a). Some DNA origami are circled (red) appearing close to the coverslip level whereas RyR2 signal is mostly in deeper tissue regions. **c**. MBM trajectories observed during the ∼40k s acquisition. **d**. Example set of localizations originating from an O1 structure after MBM correction and **e**. after additional residual drift correction. **f**. Residual drift trajectory in x determined by the correlated site-dispersion algorithm. **g**. A DNA origami structure that was oriented approximately vertically, presumably locally adhering to the tissue. Inset is the magnified area circled (red) in the overview. **h**. Overview from another dataset in which MINFLUX imaging was performed in two adjacent ROIs acquired simultaneously by scan multiplexing. ROI I contained exclusively origami (located on the coverslip where the tissue slice presumably exhibited a small crack) whereas ROI II contained almost exclusively signal from sdABs binding to RyR2 subunits. When the correlated site-dispersion algorithm was applied either to ROI I (panel **i**) or ROI II (panel **j**) the estimated residual drift trajectories look comparable suggesting that sample labeling sites can also be directly used for correction. Scale bars: a 2 µm, b 1 µm, d, e 50 nm, g 200 nm, h 5 µm, insets 10 nm.

Another dataset is shown in overview in Fig. 4h where many DNA origami structures are present in an area with a small gap in the tissue section (lower left MINFLUX ROI I) whereas a second MINFLUX ROI (ROI II, top right in Fig. 4h) mostly contains signal from RyR2s. We estimated the residual drift using the correlated site-dispersion approach either from the DNA origami ROI (ROI I) or from the RyR2 MINFLUX ROI (ROI II). The estimated residual drift trajectories were very similar (Fig. 4i and 4j). Supplementary Fig. 5 shows another sample dataset supporting this observation. This suggests that residual drift can also be directly estimated from marker sites in the absence of DNA origami signals if localization grouping into distinct repeatedly visited sites is possible.

## Discussion

We have demonstrated that DNA origami with repeat domain sites is well-suited to characterize the long-term acquisition properties of 3D MINFLUX microscopy. Use of repeat DNA-PAINT is crucial to achieving many repeat visits at DNA origami sites, which in turn enables estimating residual drift using the correlated site-dispersion approach. The methodology is also compatible with the imaging of biological samples. When DNA origami structures are included with biological samples during MINFLUX imaging one can quality assure acquisitions by detecting and correcting the residual drift remaining after application of other fiducial based drift correction (here the MBM system).

To achieve high effective labeling efficiency MINFLUX acquisitions typically need to run for an extended period of time (several hours even for micrometer sized ROIs), due to the entirely serial nature of current implementations^14^. When employing conventional DNA-PAINT these acquisitions are generally associated with extensive site-loss^14^, which we also observed when using DNA origami but could be effectively overcome with extended repeat domains^19^. This is convenient as it is compatible with standard salt buffers that do not contain any labile components. In addition, the use of repeat domains allows lowering imager concentration (by a factor which is approximately proportional to the repeat number^19^), which should benefit MINFLUX localization^23^. On first sight, the use of extended repeat domains (to provide redundancy and counter site-loss these repeat domains are not overlapping) may seem to incur a spatial resolution penalty. By contrast, site precision, a measure of the scatter of repeated localizations around single well-separated sites, was indistinguishable from the use of standard “short” (8-10 nt) docking domains, at least at the spatial resolution of ∼6 nm that we practically achieve after residual drift correction. This likely results from the rapid thermal fluctuations of the mostly single-stranded repeat domain strands which our previous simulations predict to be in the microsecond range^19^. These fluctuation timescales are fast compared to the duration of MINFLUX localizations (which are in the several millisecond range), such that the small effective PSF broadening^19^ leads to negligible reduction in localization precision.

Using DNA origami with repeat domain anchor strands provided evidence for residual drift while employing both sample stabilization to better than 1 nm as well as the MINFLUX beamline monitoring system (MBM). In scanning MINFLUX the beamline coupling into the microscope constitutes a second source of drift that, at the spatial resolution aimed for with MINFLUX (better than 10 nm), also plays a role. For long acquisitions, the MBM system to monitor and reduce beamline induced drift is essential (unless temperature stays approximately constant) and, as our data shows, it greatly reduces this drift. However, at least in the system used here, residual drift remained and could be estimated by a straightforward algorithm. The extent of beamline drift and ability of the MBM system to fully correct it may be different across installations and the methodology we introduce here can be used to compare between installations. In our experience the extent of beamline drift is strongly correlated with temperature changes (see also Supplementary Fig. 3) and was often small when temperature variations remained below ∼0.3 ℃. Under these conditions, residual drift was generally also small (< 2.5 nm). At this stage it is not clear why the tracking of gold nanoparticles by the MBM system does not fully correct beamline drift in practice – this can be further investigated with our methods acting as an assay. We believe that attachment issues are unlikely, as we remove diverging MBM bead trajectories and rely on at least 3-4 MBM beads showing strongly correlated trajectories. Again, evaluation across sites and installations may help reveal systematic factors.

The correlated site-dispersion approach to estimate drift relies on sufficiently well separated sites and “approximate” initial drift correction so that localizations can be assigned to individual sites by clustering methods. This is robustly achieved with the existing MBM correction and the presence of many repeatedly visited sites that are not too close, as ensured by the O1 and O2 DNA origami structures used here. We provide evidence that small markers exhibiting repeat domains for DNA-PAINT and typical biological target densities (as exhibited by the RyR2 labeling in our cardiac tissue samples) can also serve this purpose without explicit need for DNA origami reference structures as we illustrate in Fig. 4 and Supplementary Fig. 5. The essential property to enable this is evidently the repeated visit of individual sites (to provide reliable centroid estimates) which (in our hands) seemed difficult to achieve with dSTORM and conventional DNA-PAINT when using long-term MINFLUX acquisitions^14^. We therefore conclude that the routine use of small markers with repeat domain docking strands constitutes a convenient means to enable additional drift-correction using correlated site-dispersion. This provides an additional argument for using non-overlapping repeat domains for MINFLUX DNA-PAINT.

While the correlated site-dispersion drift correction approach has the advantage of not relying on known template structures per se it has a limited time response. In practice, it seems difficult to lower the time between traces (i.e. localizations from individual imager strands) much below 1 s^24^. Assuming that 20-50 coalesced localizations falling into a time window are required for robust drift estimates, temporal window sizes of 50 s or above are required. This leaves faster residual drift components (0.1 – 100 Hz) difficult to estimate using this approach and will require different strategies. In any case, using the approach presented here we routinely achieve site precisions around 2 nm in all directions over total acquisition periods of >30k s (>8h) compatible with FRC and FSC resolutions around 6 nm (Supplementary Fig. 3).

While the DNA origami structures we use here are mostly two-dimensional (nanoscale bread boards), information about vertical precision is readily accessible from the DNA origami recordings and due to random attachment to biological samples some DNA origami structures end up oriented mostly vertically (see Fig. 4), suggesting near isotropic localization in 3D MINFLUX imaging is achieved in practice.

In conclusion, we establish a versatile method to quality assure long-term MINFLUX acquisitions by using DNA origami with repeat-domain anchor strands as template imaging targets. This approach reveals and corrects residual “slow” drift with a time resolution of ∼50 s and is compatible with the MINFLUX imaging of biological samples. With this approach the site precision on the MINFLUX instrument in our laboratory is approximately 2 nm in all directions. Importantly, the methodology presented here will allow others to test their instruments and should enable a performance comparison across installations. With current practically achievable MINFLUX resolution in the commercial implementation we use here (∼6 nm), extended repeat-domains encounter no detectable resolution loss, further emphasizing extended repeat-domains as a versatile tool for MINFLUX DNA-PAINT.

## Materials and methods

### Sample preparation

#### Oligonucleotides and markers

Massive-Tag-Q anti-TagFP (clone 1H7) single-domain antibodies were purchased from Massive Photonics (München, Germany) with a custom-conjugation to the D2 sequence (Table 1). To enable the use of repeat-domain motifs, the D2 strand was functionalized with the 10x RD P1 oligonucleotide which has been utilized in previous super-resolution studies^12,19,9,14^. The imagers (P1s and P5, see Table 1) were modified with an ATTO 655 dye on their 3’ end. All oligonucleotides were HPLC-purified and ordered from Eurofins Genomics (Ebersberg, Germany).

#### DNA-origami production

DNA-origami constructs O1 were generated as previously described.^19^ Briefly, standard desalted oligonucleotides (staples) purchased from Integrated DNA Technologies (IDT, Coralville, USA) were used to create a flat 2D DNA structure derived from Rothemund Rectangular Origami (RRO)^25^. Picasso software^25^ was used to design eight staples carrying a 5’biotin modification, as well as staples bearing 3’ overhangs for DNA-PAINT experiments. The DNA-origami structures house six ‘D2’ (Table 1) anchors (3’ overhangs). The anchors were arranged as two rows of three anchors, with an interline distance of ∼45 nm and in-row distances of ∼35 or ∼40 nm as shown in Fig. 1a-iv. The assembly conditions have been previously described^19^ and staple sequences of the O1 design are shown in Supplementary Tables S1 and S2. Briefly, the scaffold, biotinylated oligos, 3’-overhang oligos, and non-modified oligos were mixed in 40 μL of Tris EDTA buffer (10 mM Tris-HCl, 1 mM disodium EDTA, pH 8.0) supplemented with 12.5 mM MgCl₂. The scaffold and biotinylated oligos were added at 10 nM, 3’-overhang oligos at 1 μM, and non-modified oligos at 100 nM. RRO assembly was carried out in a thermal cycler as follows. First, an 80°C incubation step °C was performed to eliminate secondary structures. Then, the assembly was allowed to occur by gradually decreasing the temperature from 60 °C to 4 °C for 3 hours, with a ramp rate of ∼1 °C per 190 s.

DNA-origami constructs O2 were generated using a similar procedure as previously described^26^. Briefly, the square-lattice DNA origami structure O2 of dimension 60 x 52.5 x 5 nm was designed using CaDNAno^27^, and is accessible at nanobase.org^28^. The DNA template was modified with 3 biotin strands at the bottom and 6 D2 DNA-PAINT docking strands at the top layer. In short, a 7249 bases scaffold (M13mp18, Tilibit Nanosystems) and 233 staple strands were folded in 1× TAE (Alfa Aesar, #J63931) and 12 mM MgCl_2_ (Alfa Aesar, #J61014) using a 1:10 scaffold/staples ratio and 1:100 for modified staples. Unmodified DNA sequences were purchased from Integrated DNA Technologies (IDT, Coralville, USA), and biotin-functionalized strands were purchased from Biomers GmbH (Ulm, Germany). For folding the solution was first heated up to 75 °C and then ramped down to 25 °C at a rate of 1 °C / 20min. Afterwards, the DNA origami structures were purified via gel electrophoresis (1 % agarose gel in 1xTAE, 12 mM MgCl_2_ buffer for 2.5 h at 4 V/cm). The appropriate band containing folded DNA origami was cut out, squeezed out from the gel and stored at −20 °C before use. The biotin modified and unmodified staples are listed in Supplementary Tables S3 and S4, respectively.

#### Preparation of DNA origami on coverslips

No. 1.5, 22 x 22 mm, glass coverslips (Menzel Gläser) were cleaned by immersing the coverslips in a beaker of acetone and sonicated for 15 minutes. Slips were air-dried and then subsequently immersed in isopropanol and sonicated for a further 15 minutes before being air-dried. Following cleaning, the coverslips were mounted on custom open-top Perspex chambers using a two-component silicone molding rubber (Pinkysil, Barnes Products, Moorebank, Australia). Coverslips were covered with 1 mg/mL solution of albumin, biotin labeled bovine (BSA-biotin A8549, Merck, Darmstadt, Germany) in PBS for 10 minutes at room temperature. Chambers were washed three times in PBS to remove excess BSA-biotin, before introducing a 1 mg/mL solution of NeutrAvidin Protein (31000, ThermoFisher Scientific, Waltham, USA) in PBS for 10 minutes at room temperature. Samples were then washed three times with PBS supplemented with 10 mM MgCl_2_. DNA-origami, stored in 1x TRIS EDTA buffer (TE, 10 mM Tris-HCl, 1 mM disodium EDTA, pH 8.0, Merck) supplemented with 12.5 mM MgCl_2_, was diluted to ∼0.2 nM in PBS containing 10 mM MgCl_2_ and incubated for 15 minutes at room temperature. Excess unbound DNA-origami was washed-out with three consecutive washes of the same buffer. On some slides together with structure O1 a small number of nano pillars were also added (design based on ^29^, with minor modifications). Signal from nano pillars was effectively rejected by the correlated site-dispersion algorithm and not used for drift estimates. For repeat-domain experiments, the functionalization strand 10x RD P1 was added at ∼200 nM in DNA-PAINT imaging buffer (PBS supplemented with an additional 500 mM NaCl_2_) to attach to D2 anchors (Table 1) for 10 minutes and subsequently washed extensively with imaging buffer to remove unbound repeat-domains.

#### Gold nanoparticle addition for MBM correction

Right before imaging, and to allow sample stage stabilization and MBM correction, gold nanoparticles (BBI Solutions, EM.GC150/4, Crumlin, United Kingdom) were diluted 1:1 in imaging buffer and incubated for 5 minutes on the samples followed by washes with imaging buffer. Attachment of the nanoparticles was monitored on the MINFLUX system. When jittering nanoparticles were observed, a 5-minute incubation with either poly-L-Lysine or imaging buffer supplemented with 10 mM MgCl_2_, followed by washes with imaging buffer, was performed. For each acquisition, a minimum of 5 nanoparticles adsorbed to the glass surface were selected for MBM tracking before starting the MINFLUX acquisition. Selected nanoparticles are also referred to as beads or MBM beads elsewhere in the text.

#### Cardiac tissue sections and RyR2 labeling

PA-RFP RyR2 mice that harbor PA-TagRFP inserted after T1365, within exon 31 of the mRyR2 gene on chromosome 13 as described previously^22^, were housed at the University of Bern. Adult mice were first anesthetized by intraperitoneal injection of pentobarbital (150 mg/kg of body weight). After the disappearance of the tail pinch reflex, heparin was intraperitoneally injected (300 IU). After approximately 10 minutes, the beating heart was rapidly excised, cannulated, and retrogradely perfused with Ca^2+^-free modified Tyrode solution (containing 140 mM NaCl, 5.4 mM KCl, 1.1 mM MgCl_2_, 1 mM NaH_2_PO_4_, 10 mM HEPES and 10 mM D-glucose, pH adjusted to 7.4 using NaOH) to remove blood. Once fluids ran clear, the heart was perfused with 5 ml of 2 % PFA diluted from stock (Electron Microscopy Sciences, 15714) in PBS at room temperature. After that, the heart was kept for an additional 10 minutes in 2 % PFA in PBS at room temperature. The samples were washed in PBS for 10 minutes, then transferred to 100 mM glycine in PBS for 15 minutes and washed again in PBS at 4 °C for 60 minutes. To cryoprotect the heart, it was then transferred to 10 % sucrose for 60 minutes at 4 °C, then to 20 % sucrose for a further 60 minutes at 4 °C, and finally to 30 % sucrose overnight at 4 °C. The next day, the heart was placed in embedding medium for cryosectioning (OCT compound) and fast frozen in 2-methylbutane placed into liquid nitrogen. Frozen tissue was stored at – 80 °C until required. Thin cryosections (3 µm) were obtained using an Epredia CryoStar NX50 Cryostat (ThermoFisher Scientific), adhered onto PLL-coated glass coverslips and stored at -20 °C until labeled.

Cardiac muscle sections on glass coverslips were mounted to custom-made open-top chambers, permeabilized with 0.1% Triton X-100 for 10 minutes, and then blocked with Massive Photonics (MP, Munich, Germany) antibody incubation solution for 1 hour. Sections were then incubated 1:100 with Massive-Tag-Q anti-TagFP single-domain antibodies (Massive-Photonics, Munich, Germany) harboring the D2 sequence diluted in MP antibody incubation solution for 2 hours at room temperature and functionalized with 10x repeat domains via the adapter strand (Table 1) followed by repeated washes with DNA-PAINT imaging buffer to remove excess adapter strands. In some experiments to better identify regions containing RyR2s, we applied counterstaining for RyR2s after staining with DNA-PAINT sdABs. Sections were incubated with single-domain antibody FluoTag-[X2] anti-TagFP-Atto488 (NanoTag Biotechnologies, Göttingen, Germany) diluted to 1:200 in MP antibody incubation solution for 60 minutes at room temperature.

#### Adding DNA origami to labeled tissue sections

DNA origami structures were added to coverslips with tissue sections (either before or after RyR2 labeling) and functionalized as described in the section on preparation of DNA origami on coverslips except for increased incubation periods and origami incubation concentration. Tissue sections were incubated in 1 mg/mL BSA-biotin for ∼45 minutes and washed in PBS, 1mg/mL NeutrAvidin was then added for ∼45 minutes and subsequently washed in PBS containing 10 mM MgCl_2_. DNA-origami was added at ∼1 nM for 1-2 hours in PBS with 10 mM MgCl_2_. 10x RD strands were added at ∼100 mM in DNA-PAINT imaging buffer for 10 minutes before being thoroughly washed in the imaging buffer. Adapter strands were added after DNA origami addition to the tissue sample and when using anti-TagFP-Q-D2 would attach to both the sdABs and the DNA origami anchors.

### Imaging and Analysis

#### MINFLUX 3D imaging

##### Instrumentation

MINFLUX recordings were performed on a commercial 3D MINFLUX system (Abberior Instruments, Göttingen, Germany) based on an inverted Olympus IX83 microscopy body equipped with a 100x, 1.45 NA UPlaXAPO oil-immersion objective. A Physik Instrumente (PI, Karlsruhe, Germany) nano XYZ stage and a 980 nm IR laser were used for active sample stabilization.

##### Acquisition parameters

Confocal scans to localize tissue slices were performed using the 488 nm laser line. MINFLUX acquisitions were carried out using a 642 nm excitation laser, and the emitted fluorescence signal was detected via two avalanche photodiodes (APD) with the following spectral windows: Cy5 near (650 – 685 nm), Cy5 far (685 – 720 nm). All confocal scans, MINFLUX recording, and laser alignment were performed with a pinhole diameter of 0.83AU. MINFLUX data were recorded with the Abberior laser at 642 nm and a software setting of 8-10 % in iterations 0-3 and x-fold increases as specified in the MINFLUX 3D localization sequence (Table 2), corresponding to ∼65 μW at the sample in iterations 0-3. For all acquisitions, the MINFLUX beamline monitoring (MBM) tracking module was used on user-selected gold particles surrounding the MINFLUX ROI. Each experiment aimed to have at least 5 gold particles tracked throughout the duration of the acquisition.

**Table 2.**
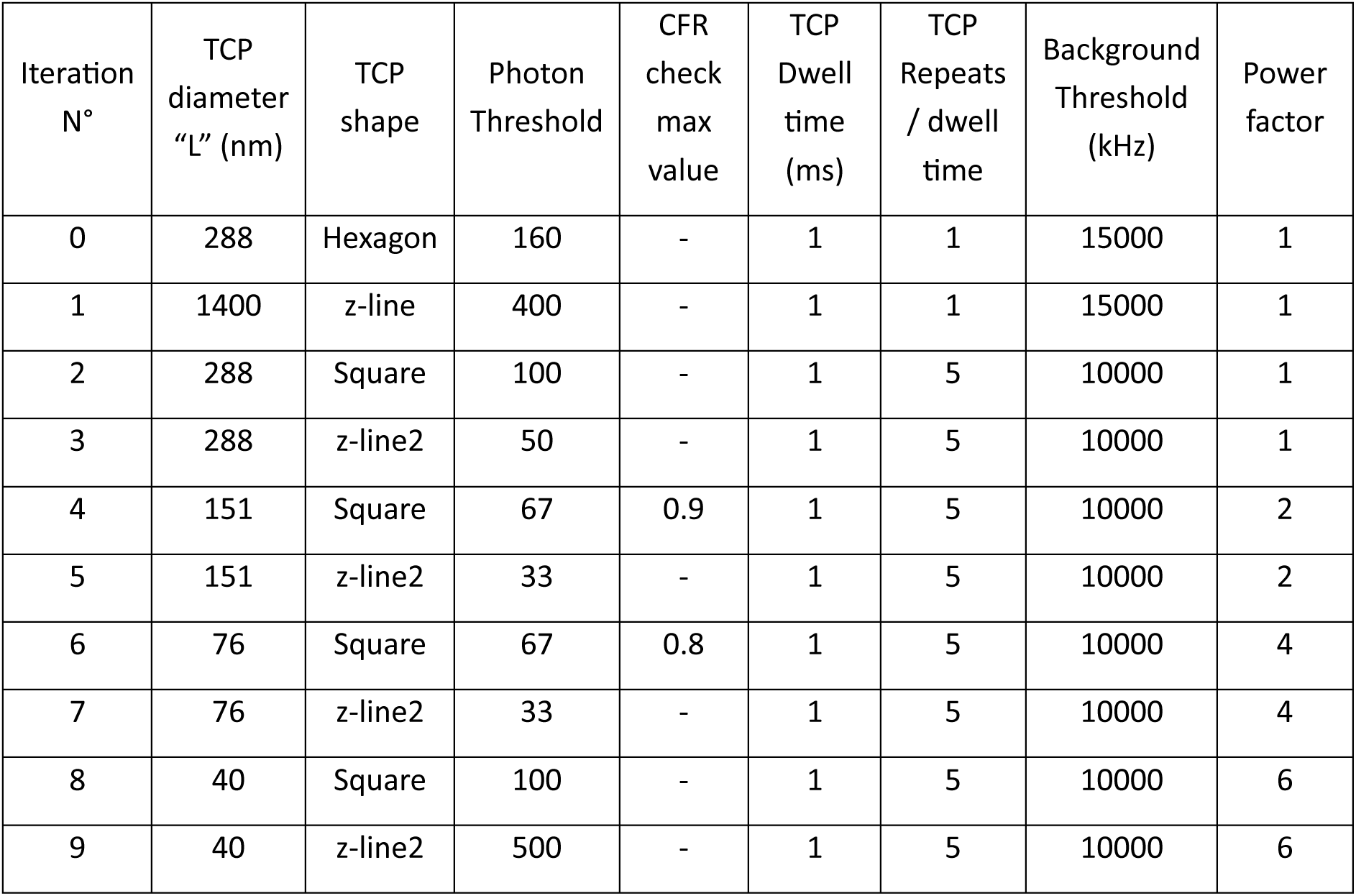
Main MINFLUX 3D Imaging sequence parameters.

##### MINFLUX localization procedure

During a 3D MINFLUX experiment, the structured excitation beam is scanned across a user-defined ROI following a hexagonal grid pattern to detect fluorescent signals. Upon detection, the iterative MINFLUX localization sequence is initiated. Initially, a defined set of coordinates, the “Target Coordinate Pattern” (TCP), is probed around the detected fluorescent signal to infer its precise localization. This TCP probing is performed at least 10 times to determine the final location. Throughout the iterative process, the TCP is recentered such that the zero-intensity center of the excitation beam coincides with the fluorophore position. Each iteration is governed by a predefined set of parameters stored in a JSON-formatted sequence file. For 3D MINFLUX imaging, the default 3D imaging sequence provided by Abberior was used (Table 2). The attributes and functions of the parameters of the sequence file were previously described^14^ (see also Supporting Fig. S2 therein).

##### Alignment and PSF verification

An Abberior nanoparticle calibration sample containing 150 nm Gold and 120 nm two-color fluorescent beads (Abberior # NP-3012) was used to evaluate the PSF shape and confirm the presence of a zero-intensity center on the 3D-shaped beam (PSF verification). PSF verification and laser alignment verification were performed before every recording using a pinhole diameter of 0.83AU as described previously^14^. Briefly, bead images were used to examine 3D donut-shaped PSF morphology and intensity homogeneity with an open pinhole. Furthermore, when necessary, the closed pinhole (0.83AU) was re-aligned to ensure the absence of clipping of the 3D PSF.

#### Software and analysis

##### Data analysis using PYME

Data exported in .npy or .zarr format from the Abberior ‘Imspector’ software were imported into the open-source Python Microscopy Environment (PYME) software^30^, http://github.com/python-microscopy/python-microscopy. Additional functionality for PYME based analysis was provided via plug-ins from the package <u>PYME-extra</u>. Zarr formatted exports generated by Imspector in DirectoryStore format were converted into zarr ZipStore format^31^ so that a single compound file was available for import. ZipStore formatted .zarr.zip files can be natively imported into PYMEVisualize with the Pyme-extra plugin package. Upon import, generally, a foreshortening factor of 0.72 was applied to all z coordinates, similar as used previously^14,32^. MBM tracks were plotted to inspect bead trajectory agreement, and divergent tracks were rejected as described in more detail in the supplementary methods. The resulting mean MBM trajectory was subtracted from the non-corrected coordinate localization as imported from the Abberior Imspector data (lnc localization data property) with the corresponding functionality implemented by the MBMcorrection module from PYME-extra. To reject low quality localizations and to effectively eliminate potential multi-emitter events from overlapping emissions, CFR and EFO filtering were conducted, as described previously. Localizations originating from the same trace as identified by the inherent ‘traceID’ property were coalesced to a single ‘coalesced localization’, as described previously^24,33^. All further analysis was generally conducted with the MBM-corrected and coalesced data. As the next step in the analysis a site-based correction algorithm was implemented. This procedure aims to identify coalesced localizations originating from the same single site, typically located on DNA-origami structures, and is described below. The complete data analysis pipeline was captured as a PYME recipe as previously described^30^ and saved as part of a session file that saves the state of the PYMEVisualize application used for display, rendering and analysis.

##### Site-based estimation of residual drift by identifying correlated site-dispersion

To estimate residual drift an estimation procedure was implemented as shown in the overview in Supplementary Fig. 2 and schematically represented in Supplementary Fig. 2a. Groups of localizations (using coalesced localizations each representing a single trace, as described above) originating from single anchor sites on DNA origami were identified by DBSCAN clustering of MBM-corrected and coalesced localizations. ε was chosen between 10 and 15 nm, depending on the separation of DNA origami sites (Supplementary Fig. 2b). ε = 15 nm was generally used for well-spaced sites of DNA origami design O1 (where sites are at least ∼35 nm apart), and ε = 10 nm for the tighter spaced sites on design O2 (see also Figs. 1a-iv, 1b-iv). The results of the DBSCAN based site grouping were also used to construct histograms of the number of site visits (see also Fig. 1). DBSCAN site groups were filtered by group size to only retain site groups containing 5 to 50 coalesced localizations (where the upper limit can be adjusted based on imaging duration and expected number of returns to individual sites). The lower limit ensures robust statistics and the upper limit rejects groups that likely represent multiple sites incorrectly “fused” by the DBCSAN clustering operation. Additional filtering to exclude “fused” or not well-isolated sites is optional, using the bounding box size of localizations in a site group or the standard deviation of group coordinates as additional filtering criteria. For each site group remaining after filtering the centroid of site localizations was calculated, as depicted in Supplementary Fig. 2c. For each localization in a site group, the shift Δs to the group centroid was determined and all shift components by coordinate axis were scatter plotted against the time localizations were obtained, as illustrated for Δx versus time in Supplementary Fig. 2d. The cloud of Δx values conveys evidence of systematic drift where correlated shifts are visible as a band of higher density of dots. The drift was estimated by determining the median shift Δx in a sliding time window (user chosen from 50 to 400 s to contain enough shift values). The sequence of median shifts through time was then smoothed (generally using locally weighted scatterplot smoothing (LOWESS)^34^ with a user selectable LOWESS fraction, typically between 5·10^-2^ and 5·10^-3^) and the resulting trajectory (green line in Supplementary Fig. 2d) provided the time course of residual drift which was in a next step removed from all coordinates. Linear interpolation was used to obtain correction values for all times at which localizations had been obtained. This estimation procedure of “correlated site-dispersion”, illustrated here for the x-axis, was equivalently conducted for y- and z-coordinates to obtain trajectories for all three axes, and the corrections were subtracted from input coordinates (i.e. coalesced localizations that had already MBM corrections applied). The corrected coordinates are shown in Supplementary Fig. 2e which visually conveys a tighter site grouping of localizations when compared with Supplementary Fig. 2c. This was quantified by calculating the standard deviation of site group coordinates, termed “site precision”, before and after residual drift correction. Site precision typically improved by 0.2-0.3 nm, sometimes more, in each coordinate axis by this procedure (see also Fig. 2f). The correction procedure was implemented in the PYME-extra package as the OrigamiSiteTracking module and the combination of site-grouping and correction can be applied in the PYMEVisualize^30^ app using the “MINFLUX>Origami>group and analyse origami sites” command. To quantify MBM correction and residual drift, trajectories were evaluated for the total correction range (maximum – minimum values) and the root mean square (RMS) deviation from no correction, i.e. RMS deviation was calculated as the square root of the mean of all squared differences. This was generally carried out for trajectories from an acquisition duration of 30k s.

##### Visualizing 3D MINFLUX data

MINFLUX localizations were visualized in the PYMEvisualize point viewer. Localizations were generally shown as OpenGL based Gaussian shaped pointsprites^30^ and colored according to one of several properties (z location or acquisition time) to obtain a Gaussian rendering in 3D. In some visualizations localizations were shown as points using the PYMEvisualize point viewer, with a typical diameter of ∼5 nm.

### Statistics and reproducibility

Data is generally given as mean or median value as stated. All p-values were obtained using the Python library package SciPy (version 1.17.0) and the ‘stats’ module by running independent t-tests using the function ‘ttest_ind’ after testing for difference of variance. A p-value <0.05 was considered as statistically significant, the label *ns* in figures denotes non-significance.

## Code availability

Processing was conducted using the PYME software package (https://github.com/python-microscopy/python-microscopy) with apps including PYMEVisualize and PYMEImage as well as by directly using PYME API functions in Jupyter notebooks (using Python 3) and the PYME plugin package PYME-extra (https://github.com/csoeller/PYME-extra). In addition, the application MBM Inspector (https://github.com/csoeller/MBM-inspector) was developed to inspect and process MBM bead trajectories.

## Acknowledgements

The authors would like to acknowledge funding support from the Swiss National Science Foundation (SNSF 310030_208109 to C. S.). The equipment was supported by the SNSF (SNSF R’Equip 316030_213543 to C. S.), the University of Bern Innovation Fund and the Faculty of Medicine. The authors would like to acknowledge the help of Dr. William T. Kaufhold for manufacturing some of the DNA origami structures used in this study.

## Author contributions

Conceptualization: C.S, A.H.C. Data curation: C.S, A.H.C, R.J., A.F.E.B. Formal analysis: C.S, A.H.C. Funding acquisition: C.S. Investigation: A.H.C, R.J., A.F.E.B Methodology: C.S, A.H.C, R.J, A.F.E.B. Project administration: C.S. Resources: R.J, A.F.E.B, K.K, G.P.A, C.S. Software: C.S, A.H.C. Supervision: C.S, A.H.C. Validation: R.J, A.H.C, A.F.E.B. Visualization: C.S, A.H.C. Writing: All authors contributed to the writing of the manuscript.

