## Supplementary Methods, Figures and Tables for "Characterizing MINFLUX imaging performance with DNA origami"

##### Processing of MBM data

MINFLUX acquisitions used the MINFLUX beamline monitoring (MBM) module[1] with user selected gold nanoparticles. Supplementary Fig. 1A shows a confocal reflection scan of a sample with MINFLUX ROI and surrounding beads selected for MBM. Beads are labeled as R<number> where <number> is a whole number generally between 0 and 20. Tracked MBM bead properties are shown during acquisition in the Imspector software and beads that are lost during tracking (either because of physical detachment but more often because the MBM module decides to drop a bead based on criteria that are not fully discernible from the software description) before the acquisition is terminated are automatically deselected for correction. A mean MBM trajectory is calculated by Abberior Imspector from all tracked beads and subtracted from the raw MINFLUX coordinates in 'real-time'. For reference, the MINFLUX data also contains the raw uncorrected coordinates which are available via the `loc_nc` (not corrected localization coordinates) property. The data in Fig. 1 shows the MBM corrected data with the MBM correction applied during acquisition whereas all other data shown (Fig. 2 and later) applied the MBM correction by extracting the MBM bead trajectories from the exported data, processing the individual MBM bead data and applying MBM corrections from within the PYMEVizualize localization processing application. Moving the MBM processing and drift correction into our data analysis pipeline allowed more control and inspection of the MBM data.

###### *Extraction of MBM data*

The full MBM information was extracted as individual bead trajectories from the ZARR file (present in the zarr array grd/mbm/points) exported from Abberior Imspector. We developed a graphical editor (MBM Inspector) displaying MBM bead trajectories (Screenshots in Supplementary Fig. 1b). This allowed identifying bead trajectories diverging from well co-aligned trajectories. These trajectories were manually deselected and from the remaining trajectories a mean MBM trajectory was generated. Diverging trajectories were present in most datasets and could not be unequivocally related to protocol or nanoparticle attachment issues. This observation also motivated selecting a sizable number of beads (typically around 10) for MBM tracking so that correlated bead movements could be distinguished from outliers that were removed in the manner described above. In Supplementary Fig. 1b deselected bead trajectories are shown translucent, and the diverging trajectories are discernible in this way. When inspection is completed, the bead selection information is saved as a JSON file which notes selected and deselected MBM beads identified by their R<number> labels.

##### *Applying MBM corrections in the PYMEVisualize analysis pipeline*

The saved bead selection information (in JSON format) together with the individual MBM bead trajectories is then read by the `MBMcorrection` module that is part of PYME-extra. Within the `MBMcorrection` module, the mean MBM trajectory for correction is calculated only from selected beads, ignoring deselected beads, and generating a mean trajectory with optional smoothing applied. The mean MBM trajectory is then used in the localization processing pipeline within PYMEVizualize to correct the raw uncorrected localizations (i.e. coordinates extracted from the `loc_nc` property). An example MBM trajectory is shown in Supplementary Fig. 1c which also shows comparison curves that show the total drift trajectory taking additionally the residual drift into account which was determined by the correlated site-dispersion algorithm. In this example, the deviations identified as residual drift were relatively small, i.e. the residual drift did not exceed ~2.5 nm in either direction (Supplementary Fig. 2d).

Processing the MBM bead information in this way allowed greater control, particularly identification of diverging bead trajectories and exclusion of those bead trajectories from calculation of the mean MBM drift trajectory. In general, at least 3-4 MBM beads, often more, showed good trajectory agreement increasing confidence in the quality of the MBM correction.

The correlated site-dispersion algorithm (see main text methods) used coalesced coordinates that were MBM corrected within our analysis pipeline as described above as input data to estimate residual drift trajectories.

#### Fourier Ring and Shell Correlation

To estimate spatial image resolution we calculated the Fourier Ring Correlation (FRC)[2,3] or the Fourier Shell Correlation (FSC). To obtain FRC or FSC values, raw MINFLUX localizations were grouped into two separate groups using the "Split by Time Blocks for FRC" functionality of PYMEVisualize using a time block size of 30 s. For FRC measurements the two groups of localizations were rendered as 2D images using the Gaussian rendering method of PYMEVisualize and a pixel size of 3 nm. FRC resolution was determined as the inverse of the 1/7 threshold frequency using the FRC module of `PYME-extra`. For FSC calculations, two groups of localizations were rendered into two 3D volume images using the Gaussian rendering method of PYMEVisualize and a pixel size of 3 nm. The volume images were saved via the PYME-extra menu function '`save MRC volumes for FSC`' in MRC format using the Python package `mrcfile` which provides a Python implementation of the MRC2014 file format. The data was then uploaded to the EMDB FSC server at https://www.ebi.ac.uk/emdb/validation/fsc which calculated the FSC curve from the provided data. The FSC resolution was determined as the inverse of the 1/7 threshold spatial frequency.

### Supplementary Tables

#### DNA Origami Designs

**Table S1:** DNA sequences for DNA origami synthesis O1 – sequences with 5' biotin end modifications (5' → 3').

| |
|---|
| /5Biosg/ATTAAGTTTACCGAGCTCGAATTCGGGAAACCTGTCGTGC |
| /5Biosg/ATAAGGGAACCGGATATTCATTACGTCAGGACGTTGGGAA |
| /5Biosg/GCGATCGGCAATTCCACACAACAGGTGCCTAATGAGTG |
| /5Biosg/TTGTGTCGTGACGAGAAACACCAAATTTCAACTTTAAT |
| /5Biosg/ATTCATTTTTGTTTGGATTATACTAAGAAACCACCAGAAG |
| /5Biosg/CACCCTCAGAAACCATCGATAGCATTGAGCCATTTGGGAA |
| /5Biosg/AACAATAACGTAAAACAGAAATAAAAATCCTTTGCCCGAA |
| /5Biosg/AGCCACCACTGTAGCGCGTTTTCAAGGGAGGGAAGGTAAA |

**Table S2:** DNA sequences for DNA origami synthesis O1 – sequences without biotin modifications (5' → 3').

| Oligo Name | Sequence | Overhang Set |
|---|---|---|
| 21[32]23[31]D1 | TTTTCACTCAAAGGGCGAAAAACCATCACC | |
| 19[32]21[31]BLK | GTCGACTTCGGCCAACGCGCGGGGTTTTTC **TTTTAGGTAAATTTTGATTGTGAGGAAG** | **D2** |
| 17[32]19[31]BLK | TGCATCTTTCCCAGTCACGACGGCCTGCAG | |
| 15[32]17[31]BLK | TAATCAGCGGATTGACCGTAATCGTAACCG | |
| 13[32]15[31]BLK | AACGCAAAATCGATGAACGGTACCGGTTGA | |
| 11[32]13[31]BLK | AACAGTTTTGTACCAAAAACATTTTATTTC | |
| 9[32]11[31]BLK | TTTACCCCAACATGTTTTAAATTTCCATAT | |
| 7[32]9[31]BLK | TTTAGGACAAATGCTTTAAACAATCAGGTC | |
| 5[32]7[31]BLK | CATCAAGTAAAACGAACTAACGAGTTGAGA | |
| 3[32]5[31]BLK | AATACGTTTGAAAGAGGACAGACTGACCTT | |

| | | |
|---|---|---|
| 1[32]3[31]BLK | AGGCTCCAGAGGCTTTGAGGACACGGGTAA **TTTTAGGTAAATTTTGATGTGAGGAAG** | **D2** |
| 0[47]1[31]D1 | AGAAAGGAACAACTAAAGGAATTCAAAAAAA | |
| 23[32]22[48]BLK | CAAATCAAGTTTTTTGGGGTCGAAACGTGGA | |
| 22[47]20[48]BLK | CTCCAACGCAGTGAGACGGGCAACCAGCTGCA | |
| 20[47]18[48]BLK | TTAATGAACTAGAGGATCCCCGGGGGGTAACG | |
| 18[47]16[48]BLK | CCAGGGTTGCCAGTTTGAGGGGACCCGTGGGA | |
| 16[47]14[48]BLK | ACAAACGGAAAAGCCCCAAAAACACTGGAGCA | |
| 14[47]12[48]BLK | AACAAGAGGGATAAAAATTTTTAGCATAAAGC | |
| 12[47]10[48]BLK | TAAATCGGGATTCCCAATTCTGCGATATAATG | |
| 10[47]8[48]BLK | CTGTAGCTTGACTATTATAGTCAGTTCATTGA | |
| 8[47]6[48]BLK | ATCCCCCTATACCACATTCAACTAGAAAAATC | |
| 6[47]4[48]BLK | TACGTTAAAGTAATCTTGACAAGAACCGAACT | |
| 4[47]2[48]BLK | GACCAACTAATGCCACTACGAAGGGGGTAGCA | |
| 2[47]0[48]BLK | ACGGCTACAAAAGGAGCCTTTAATGTGAGAAT | |
| 21[56]23[63]BLK | AGCTGATTGCCCTTCAGAGTCCACTATTAAAGGGTGCCGT | |
| 15[64]18[64]BLK | GTATAAGCCAACCCGTCGGATTCTGACGACAGTATCGGCCGCAAG GCG | |
| 13[64]15[63]BLK | TATATTTTGTCATTGCCTGAGAGTGGAAGATT | |
| 11[64]13[63]BLK | GATTTAGTCAATAAAGCCTCAGAGAACCCTCA | |
| 9[64]11[63]BLK | CGGATTGCAGAGCTTAATTGCTGAAACGAGTA | |
| 7[56]9[63]BLK | ATGCAGATACATAACGGGAATCGTCATAAATAAAGCAAAG | |
| 1[64]4[64]BLK | TTTATCAGGACAGCATCGGAACGACACCAACCTAAAACGAGGTCA ATC | |
| 0[79]1[63]BLK | ACAACTTTCAACAGTTTCAGCGGATGTATCGG | |
| 23[64]22[80]BLK | AAAGCACTAAATCGGAACCCTAATCCAGTT | |
| 22[79]20[80]BLK | TGGAACAACCGCCTGGCCCTGAGGCCCGCT | |

| | |
|---|---|
| 20[79]18[80]BLK | TTCCAGTCGTAATCATGGTCATAAAAGGGG |
| 18[79]16[80]BLK | GATGTGCTTCAGGAAGATCGCACAATGTGA |
| 16[79]14[80]BLK | GCGAGTAAAAATATTTAAATTGTTACAAAG |
| 14[79]12[80]BLK | GCTATCAGAAATGCAATGCCTGAATTAGCA |
| 12[79]10[80]BLK | AAATTAAGTTGACCATTAGATACTTTTGCG |
| 10[79]8[80]BLK | GATGGCTTATCAAAAAGATTAAGAGCGTCC |
| 8[79]6[80]BLK | AATACTGCCCAAAAGGAATTACGTGGCTCA |
| 6[79]4[80]BLK | TTATACCACCAAATCAACGTAACGAACGAG |
| 4[79]2[80]BLK | GCGCAGACAAGAGGCAAAAGAATCCCTCAG |
| 2[79]0[80]BLK | CAGCGAAACTTGCTTTCGAGGTGTTGCTAA |
| 21[96]23[95]BLK | AGCAAGCGTAGGGTTGAGTGTTGTAGGGAGCC |
| 19[96]21[95]BLK | CTGTGTGATTGCGTTGCGCTCACTAGAGTTGC |
| 17[96]19[95]BLK | GCTTTCCGATTACGCCAGCTGGCGGCTGTTTC |
| 15[96]17[95]BLK | ATATTTTGGCTTTCATCAACATTATCCAGCCA |
| 13[96]15[95]BLK | TAGGTAAACTATTTTTGAGAGATCAAACGTTA |
| 11[96]13[95]BLK | AATGGTCAACAGGCAAGGCAAAGAGTAATGTG |
| 9[96]11[95]BLK | CGAAAGACTTTGATAAGAGGTCATATTTCGCA |
| 7[96]9[95]BLK | TAAGAGCAAATGTTTAGACTGGATAGGAAGCC |
| 5[96]7[95]BLK | TCATTCAGATGCGATTTTAAGAACAGGCATAG |
| 3[96]5[95]BLK | ACACTCATCCATGTTACTTAGCCGAAAGCTGC |
| 1[96]3[95]BLK | AAACAGCTTTTTGCGGGATCGTCAACACTAAA |
| 0[111]1[95]BLK | TAAATGAATTTTCTGTATGGGATTAATTTCTT |
| 23[96]22[112]BLK | CCCGATTTAGAGCTTGACGGGGAAAAAGAATA |
| 22[111]20[112]BLK | GCCCGAGAGTCCACGCTGGTTTGCAGCTAACT |
| 20[111]18[112]BLK | CACATTAAAATTGTTATCCGCTCATGCGGGCC |
| 18[111]16[112]BLK | TCTTCGCTGCACCGCTTCTGGTGCGGCCTTCC |
| 16[111]14[112]BLK | TGTAGCCATTAAAATTCGCATTAAATGCCGGA |
| 14[111]12[112]BLK | GAGGGTAGGATTCAAAAGGGTGAGACATCCAA |
| 12[111]10[112]BLK | TAAATCATATAACCTGTTTAGCTAACCTTTAA |
| 10[111]8[112]BLK | TTGCTCCTTTCAAATATCGCGTTTGAGGGGGT |

| | | |
|---|---|---|
| 8[111]6[112]BLK | AATAGTAAACACTATCATAACCCTCATTGTGA | |
| 6[111]4[112]BLK | ATTACCTTTGAATAAGGCTTGCCCAAATCCGC | |
| 4[111]2[112]BLK | GACCTGCTCTTTGACCCCCAGCGAGGGAGTTA | |
| 2[111]0[112]BLK | AAGGCCGCTGATACCGATAGTTGCGACGTTAG | |
| 21[120]23[127]BLK | CCCAGCAGGCGAAAAATCCCTTATAAATCAAGCCGGCG | |
| 15[128]18[128]BLK | TAAATCAAAATAATTCGCGTCTCGGAAACCAGGCAAAGGGAAGG | |
| 13[128]15[127]BLK | GAGACAGCTAGCTGATAAATTAATTTTTGT | |
| 11[128]13[127]BLK | TTTGGGGATAGTAGTAGCATTAAAAGGCCG | |
| 9[128]11[127]BLK | GCTTCAATCAGGATTAGAGAGTTATTTTCA | |
| 7[120]9[127]BLK | CGTTTACCAGACGACAAAGAAGTTTTGCCATAATTCGA | |
| 1[128]4[128]BLK | TGACAACTCGCTGAGGCTTGCATTATACCAAGCGCGATGATAAA | |
| 0[143]1[127]BLK | TCTAAAGTTTTGTCGTCTTTCCAGCCGACAA | |
| 21[160]22[144]BLK | TCAATATCGAACCTCAAATATCAATTCCGAAA | |
| 19[160]20[144]BLK | GCAATTCACATATTCCTGATTATCAAAGTGTA | |
| 17[160]18[144]BLK | AGAAAACAAAGAAGATGATGAAACAGGCTGCG | |
| 15[160]16[144]BLK | ATCGCAAGTATGTAAATGCTGATGATAGGAAC | |
| 13[160]14[144]BLK | GTAATAAGTTAGGCAGAGGCATTTATGATATT | |
| 11[160]12[144]BLK | CCAATAGCTCATCGTAGGAATCATGGCATCAA | |
| 9[160]10[144]BLK | AGAGAGAAAAAAATGAAAATAGCAAGCAAACT | |
| 7[160]8[144]BLK | TTATTACGAAGAACTGGCATGATTGCGAGAGG | |
| 5[160]6[144]BLK | GCAAGGCCTCACCAGTAGCACCATGGGCTTGA | |
| 3[160]4[144]BLK | TTGACAGGCCACCACCAGAGCCGCGATTTGTA | |
| 1[160]2[144]BLK | TTAGGATTGGCTGAGACTCCTCAATAACCGAT | |
| 0[175]0[144]BLK | TCCACAGACAGCCCTCATAGTTAGCGTAACGA | |
| 23[128]23[159]D1 | AACGTGGCGAGAAAGGAAGGGAAACCAGTAA | |
| 22[143]21[159]BLK | TCGGCAAATCCTGTTTGATGGTGGACCCTCAA<br>**TTTTAGGTAAATTTTGATTGTGAGGAAG** | **D2** |

| | | |
|---|---|---|
| 20[143]19[159]BLK | AAGCCTGGTACGAGCCGGAAGCATAGATGATG | |
| 18[143]17[159]BLK | CAACTGTTGCGCCATTCGCCATTCAAACATCA | |
| 16[143]15[159]BLK | GCCATCAAGCTCATTTTTTAACCACAAATCCA | |
| 14[143]13[159]BLK | CAACCGTTTCAAATCACCATCAATTCGAGCCA | |
| 12[143]11[159]BLK | TTCTACTACGCGAGCTGAAAAGGTTACCGCGC | |
| 10[143]9[159]BLK | CCAACAGGAGCGAACCAGACCGGAGCCTTTAC | |
| 8[143]7[159]BLK | CTTTTGCAGATAAAAACCAAAATAAAGACTCC | |
| 6[143]5[159]BLK | GATGGTTTGAACGAGTAGTAAATTTACCATTA | |
| 4[143]3[159]BLK | TCATCGCCAACAAAGTACAACGGACGCCAGCA **TTTTAGGTAAATTTTGATTGTGAGGAAG** | **D2** |
| 2[143]1[159]D1 | ATATTCGGAACCATCGCCCACGCAGAGAAGGA | |
| 23[160]22[176]BLK | TAAAAGGGACATTCTGGCCAACAAAGCATC | |
| 22[175]20[176]BLK | ACCTTGCTTGGTCAGTTGGCAAAGAGCGGA | |
| 20[175]18[176]BLK | ATTATCATTCAATATAATCCTGACAATTAC | |
| 18[175]16[176]BLK | CTGAGCAAAAATTAATTACATTTTGGGTTA | |
| 16[175]14[176]BLK | TATAACTAACAAAGAACGCGAGAACGCCAA | |
| 14[175]12[176]BLK | CATGTAATAGAATATAAAGTACCAAGCCGT | |
| 12[175]10[176]BLK | TTTTATTTAAGCAAATCAGATATTTTTTGT | |
| 10[175]8[176]BLK | TTAACGTCTAACATAAAAACAGGTAACGGA | |
| 8[175]6[176]BLK | ATACCCAACAGTATGTTAGCAAATTAGAGC | |
| 6[175]4[176]BLK | CAGCAAAAGGAAACGTCACCAATGAGCCGC | |
| 4[175]2[176]BLK | CACCAGAAAGGTTGAGGCAGGTCATGAAAG | |
| 2[175]0[176]BLK | TATTAAGAAGCGGGGTTTTGCTCGTAGCAT | |
| 21[184]23[191]BLK | TCAACAGTTGAAAGGAGCAAATGAAAAATCTAGAGATAGA | |
| 15[192]18[192]BLK | TCAAATATAACCTCCGGCTTAGGTAACAATTTCATTTGAAGGCGAATT | |
| 13[192]15[191]BLK | GTAAAGTAATCGCCATATTTAACAAAACTTTT | |
| 11[192]13[191]BLK | TATCCGGTCTCATCGAGAACAAGCGACAAAAG | |
| 9[192]11[191]BLK | TTAGACGGCCAAATAAGAAACGATAGAAGGCT | |

| | |
|---|---|
| 7[184]9[191]BLK | CGTAGAAAATACATACCGAGGAAACGCAATAAGAAGCGCA |
| 1[192]4[192]BLK | GCGGATAACCTATTATTCTGAAACAGACGATTGGCCTTGAAGAGCCAC |
| 0[207]1[191]BLK | TCACCAGTACAAACTACAACGCCTAGTACCAG |
| 23[192]22[208]BLK | ACCCTTCTGACCTGAAAGCGTAAGACGCTGAG |
| 22[207]20[208]BLK | AGCCAGCAATTGAGGAAGGTTATCATCATTTT |
| 20[207]18[208]BLK | GCGGAACATCTGAATAATGGAAGGTACAAAAT |
| 18[207]16[208]BLK | CGCGCAGATTACCTTTTTTAATGGGAGAGACT |
| 16[207]14[208]BLK | ACCTTTTTATTTTAGTTAATTTCATAGGGCTT |
| 14[207]12[208]BLK | AATTGAGAATTCTGTCCAGACGACTAAACCAA |
| 12[207]10[208]BLK | GTACCGCAATTCTAAGAACGCGAGTATTATTT |
| 10[207]8[208]BLK | ATCCCAATGAGAATTAACTGAACAGTTACCAG |
| 8[207]6[208]BLK | AAGGAAACATAAAGGTGGCAACATTATCACCG |
| 6[207]4[208]BLK | TCACCGACGCACCGTAATCAGTAGCAGAACCG |
| 4[207]2[208]BLK | CCACCCTCTATTCACAAACAAATACCTGCCTA |
| 2[207]0[208]BLK | TTTCGGAAGTGCCGTCGAGAGGGTGAGTTTCG |
| 21[224]23[223]BLK | CTTTAGGGCCTGCAACAGTGCCAATACGTG |
| 19[224]21[223]BLK | CTACCATAGTTTGAGTAACATTTAAAATAT |
| 17[224]19[223]BLK | CATAAATCTTTGAATACCAAGTGTTAGAAC |
| 15[224]17[223]BLK | CCTAAATCAAAATCATAGGTCTAAACAGTA |
| 13[224]15[223]BLK | ACAACATGCCAACGCTCAACAGTCTTCTGA |
| 11[224]13[223]BLK | GCGAACCTCCAAGAACGGGTATGACAATAA |
| 9[224]11[223]BLK | AAAGTCACAAAATAAACAGCCAGCGTTTTA |
| 7[224]9[223]BLK | AACGCAAAGATAGCCGAACAAACCCTGAAC |
| 5[224]7[223]BLK | TCAAGTTTCATTAAAGGTGAATATAAAAGA |
| 3[224]5[223]BLK | TTAAAGCCAGAGCCGCCACCCTCGACAGAA |
| 1[224]3[223]BLK | GTATAGCAAACAGTTAATGCCCAATCCTCA |
| 0[239]1[223]BLK | AGGAACCCATGTACCGTAACACTTGATATAA |
| 23[224]22[240]BLK | GCACAGACAATATTTTTGAATGGGGTCAGTA |

| | | |
|---|---|---|
| 22[239]20[240]BLK | TTAACACCAGCACTAACAACTAATCGTTATTA | |
| 20[239]18[240]BLK | ATTTTAAAATCAAAATTATTTGCACGGATTCG | |
| 18[239]16[240]BLK | CCTGATTGCAATATATGTGAGTGATCAATAGT | |
| 16[239]14[240]BLK | GAATTTATTTAATGGTTTGAAATATTCTTACC | |
| 14[239]12[240]BLK | AGTATAAAGTTCAGCTAATGCAGATGTCTTTC | |
| 12[239]10[240]BLK | CTTATCATTCCCGACTTGCGGGAGCCTAATTT | |
| 10[239]8[240]BLK | GCCAGTTAGAGGGTAATTGAGCGCTTTAAGAA | |
| 8[239]6[240]BLK | AAGTAAGCAGACACCACGGAATAATATTGACG | |
| 6[239]4[240]BLK | GAAATTATTGCCTTTAGCGTCAGACCGGAACC | |
| 4[239]2[240]BLK | GCCTCCCTCAGAATGGAAAGCGCAGTAACAGT | |
| 2[239]0[240]BLK | GCCCGTATCCGGAATAGGTGTATCAGCCCAAT | |
| 21[248]23[255]BLK | AGATTAGAGCCGTCAAAAAACAGAGGTGAGGCCTATTAGT | |
| 15[256]18[256]BLK | GTGATAAAAAGACGCTGAGAAGAGATAACCTTGCTTCTGTTCGGG AGA | |
| 13[256]15[255]BLK | GTTTATCAATATGCGTTATACAAACCGACCGT | |
| 11[256]13[255]BLK | GCCTTAAACCAATCAATAATCGGCACGCGCCT | |
| 9[256]11[255]BLK | GAGAGATAGAGCGTCTTTCCAGAGGTTTTGAA | |
| 7[248]9[255]BLK | GTTTATTTTGTCACAATCTTACCGAAGCCCTTTAATATCA | |
| 1[256]4[256]BLK | CAGGAGGTGGGGTCAGTGCCTTGAGTCTCTGAATTTACCGGGAA CCAG | |
| 0[271]1[255]BLK | CCACCCTCATTTTCAGGGATAGCAACCGTACT | |
| 23[256]22[272]D1 | CTTTAATGCGCGAACTGATAGCCCCACCAG | |
| 22[271]20[272]BLK | CAGAAGATTAGATAATACATTTGTCGACAA **TTTTAGGTAAATTTTGATTGTGAGGAAG** | **D2** |
| 20[271]18[272]BLK | CTCGTATTAGAAATTGCGTAGATACAGTAC | |
| 18[271]16[272]BLK | CTTTTACAAAATCGTCGCTATTAGCGATAG | |
| 16[271]14[272]BLK | CTTAGATTTAAGGCGTTAAATAAAGCCTGT | |

| | | |
|---|---|---|
| 14[271]12[272]BLK | TTAGTATCACAATAGATAAGTCCACGAGCA | |
| 12[271]10[272]BLK | TGTAGAAATCAAGATTAGTTGCTCTTACCA | |
| 10[271]8[272]BLK | ACGCTAACACCCACAAGAATTGAAAATAGC | |
| 8[271]6[272]BLK | AATAGCTATCAATAGAAAATTCAACATTCA | |
| 6[271]4[272]BLK | ACCGATTGTCGGCATTTTCGGTCATAATCA | |
| 4[271]2[272]BLK | AAATCACCTTCCAGTAAGCGTCAGTAATAA **TTTTAGGTAAATTTTGATTGTGAGGAAG** | **D2** |
| 2[271]0[272]D1 | GTTTTAACTTAGTACCGCCACCCAGAGCCA | |

Table S3. DNA sequences for DNA origami synthesis O2 – modified sequences with 5’ biotin end modifications (5’ → 3’).

| |
|---|
| /5Biotin/TTGGAGAGGCGCAAGCGGAATCGGCAAAATCCCT |
| /5Biotin/TTCGCCTCCCTCAGAGCCAGCGTTTGGTAATCAG |
| /5Biotin/TTATGAAAGTTATAGCCCCCTCAGACATTCCAC |

Table S4. DNA sequences for DNA origami synthesis O2 – sequences without biotin modifications (5’ → 3’).

| **OLIGO NAME** | **SEQUENCE** | **Overhang Set** |
|---|---|---|
| 18[119] 16[104] Core | AAGATGATGCTTTGAAAGTAGTAGCATTAACA | |
| 30[55] 28[40] Core | TTTGCCAGTAAATCAAAAATCAGATATAGAAG | |
| 6[55] 4[40] Core | CCATCACGATTAAAGGTTTGACGAGCACGTAT | |
| 18[55] 16[40] Core | CCTTTTTTCAGATGAAAATGGAAGGGTTAGAA | |
| 6[119] 4[104] Core | TTGTAAAAGGTCGACTTAAAGCCTGGGGTGCC | |
| 30[119] 28[104] Core | CCTAAAACATCTTTGACCGAACTGACCAACTT | |
| 23[120] 21[135] Core | GAAGTTTTAAATATCGCGAGAAAACTTTTTCA | |
| 22[119] 20[104] Core | GTTTGAAAAAAGAACGCGTTTTAATTCGAGCT | |
| 9[168] 12[168] Core | AATGGGATTTGTTAAATTTAAATTATCTACAA | |
| 20[39] 25[39] Core | AGATTAAACGCTCATTTAGTAAAGGTAA | |
| 2[87] 0[72] Core | GGGAGCCCAAGCACTATTGGAACAAGAGTCCA | |
| 25[4] 24[4] Core | CCCCTAAGAGAATATAAAGTATTTTCGAGCCAGTAACCCC | |
| 23[56] 21[71] Core | GAGAATCGTCCTTGAATTAGGTTGGGTTATAT | |

| | |
|---|---|
| 26[55] 23[55] Core | TAATATCCTGTCCAGAAACAACGCGCTTAATT |
| 37[40] 40[40] Core | AAATATTGTAGCAAGGGCAGCACCCCATCTTT |
| 3[56] 1[71] Core | AAGTGTAGGAACGTGGGTTTTTTGGGGTCGAG |
| 13[72] 16[72] Core | TAGATAATCAAACAATCTGATTATATTGTTTG |
| 16[192] 15[198] Core | CCCCTCAATAACCTGTTAACATTATGACCCTGTAATCCCC |
| 6[198] 8[168] Core | CCCCACGCCAGCTGGCGAAAGGGGGATGTGCGCGATCGG TCGTAACC |
| 13[168] 16[168] Core | AATTAATGTTATTTCAGTACCAAATAGCTATA |
| 16[39] 21[39] Core | CCTACCAATCGTCGCAGTACATAGACTAC |
| 3[20] 0[15] Core | CCCCAAGCGAAAACCGTCTATCAGGCCCC |
| 5[40] 8[40] Core | GGAGGCCGCAAATTAAAGAACTCAATCGTCTG |
| 17[104] 19[119] Core | GCCTGATTGAAACAAAGTCATTTTTGCGGATG |
| 37[104] 40[104] Core | GTACCAGGTAGGATTAGATACAGGAAGCGTCA |
| 5[136] 8[136] Core | TACCGAGCGCCAGGGTCGCCATTCCGACGACA |
| 8[167] 13[167] Core | GTGCATCTGCAAATATCAGCTCAGCTGATA |
| 31[9] 28[28] Core | CCCCCCAAATAAGAAACGATTTTCGGT |
| 24[27] 22[9] Core | AGGCACCGACAATCATATGCGTTATACAAATCCCC |
| 9[4] 8[4] Core | CCCCGTCACACGACCAGTAAATTGGCAGATTCACCACCCC |
| 28[167] 33[167] Core | CATGTTACTCGGAACGTTTCCATTTAATTGT |
| 4[182] 9[167] Core | CCCCTGTGAAATTGTTATCCGCGGGAAGGTGCAAGGCTTG ACCGT |
| 25[72] 28[72] Core | CATGTTCATAGATAAGTCGAGAACTCATTACC |
| 40[39] 40[15] Core | TCATAATCAAAATCACCGGACCCC |
| 36[192] 35[198] Core | CCCCCCGCCACCCTCAGACGATCTAAAGTTTTGTCGCCCC |
| 12[39] 17[39] Core | TGGTCAGAAAGAAAGTTATTATTTCAGG |
| 15[88] 14[88] Core | GGCAATTCATCATACGTAAAGATAAGTATT |
| 24[71] 29[71] Core | CTATCATACGCACTCATCCTGAACGCTATTTT |
| 4[71] 9[71] Core | CGCGTACTGGTAATATGAGTAAAAGACCTGAA |
| 11[88] 10[88] Core | GCAAATGAATGTCAACCAGCTTTAATATTT |
| 4[135] 9[135] Core | CCGGAAGCGCGCCATTTTTCCCAGCGTCGGAT |
| 24[192] 23[198] Core | CCCCCGGAACAACATTACTGCGGAATCGTCATAAATCCCC |

| | |
|---|---|
| 40[182] 38[152] Core | CCCCCACAAACAAATAAATCCTCATTAAAGCAGGTCATCTG AAAC |
| 17[168] 20[168] Core | TTAGATACCAGTTGATAAATATGCCAAAGCGG |
| 9[40] 12[40] Core | GGCCAACATAAAACATCGGTCAGTTCAATATC |
| 27[120] 25[135] Core | AGGCGCATAACTAATGTTAAGAACTGGCTCAT |
| 39[20] 36[15] Core | CCCCATGAAACCGGCGACATTCAACCCC |
| 41[136] 39[151] Core | TGACAGGAGGTTGAGGCCAGAATGTGCCTTGA |
| 5[104] 8[104] Core | TGCCTGCACGACGGCCTTCTGGTGTCCAGCCA |
| 18[198] 17[192] Core | CCCCGAAGTTTCATTCCATATAAATTTCGCAAATGGCCCC |
| 27[152] 22[152] Core | TAATCTTGTCATCAGTTGGGAACAGAAAAC |
| 39[120] 34[120] Core | GGTAATAATCAGGGATTGCCGTCGGCGGAGTG |
| 12[192] 11[198] Core | CCCCCTATTTTTGAGAGGTAAACGTTAATATTTTGTCCCC |
| 28[135] 33[135] Core | GTCAATCAGGGATCGTACTACGAAATCTCCAA |
| 10[151] 7[151] Core | ACCAATAGGGAACAAATGAGGGGAAGGCTGCG |
| 32[27] 30[9] Core | AGAGAATAAGAGCCATATTATTTATCCCAATCCCC |
| 33[136] 36[136] Core | AAAAAAGGACTTTCAAGCCTGTAGGCCACCAC |
| 31[152] 26[152] Core | AGACAGCATTAGCCGTACAACGCTGACGAG |
| 20[192] 19[198] Core | CCCCTTATAGTCAGAAGAACTAAAGTACGGTGTCTGCCCC |
| 17[4] 16[4] Core | CCCCCAGAAATAAAGAAATTTTATTTGCACGTAAAACCCC |
| 20[167] 25[167] Core | ATTGCATCGGATAGCAAACAGTTGAAAAATC |
| 19[9] 16[28] Core | CCCCAATAACCTTGCTTCTGTAATATC |
| 34[151] 31[151] Core | GCTAAACACTCCAAAATGCGCCGAGCAGCGAA |
| 12[135] 17[135] Core | GCAAACAATTAAGCAATATTTTAATACAAAAT |
| 21[72] 23[87] Core | AACTATATAAGAATAATTACCAGACGACGATA |
| 35[56] 30[56] Core | AAGGTGGCTGAACAAAGCTATCTTGAGCCTAA |
| 38[119] 35[119] Core | AGAAGGATCGGATAAGAGCAAGCCTTCGTCAC |
| 9[104] 12[104] Core | ATTAAATGTCCTGTAGTCATATGTTAATCGTA |
| 15[152] 10[152] Core | AAAGCTAAAGGTCATCCGTTCTATTTTTTA |
| 28[27] 26[9] Core | ATTCACTTGCGGCATGTAGAAACCAATCAATCCCC |
| 32[167] 37[167] Core | CTTGATACCTCATAGTCTGTATGTGTATCA |
| 37[15] 34[9] Core | CCCCCCGATTGAGGGACTGGCATGATTAAGACTCCCCCC |

| | |
|---|---|
| 14[198] 13[192] Core | CCCCACTTTTGCGGGAGAAGCCTCCGGAGAGGGTAGCCCC |
| 10[55] 7[55] Core | GATAGCCCGAGATAGAGACGCTCAAACTATCG |
| 10[119] 7[119] Core | TCTGGCCTTGAGCGAGGATCGCACCCGGAAAC |
| 7[88] 6[88] Core | GCCAGCCAGCACCGCAGTGCCAATTTTTAT |
| 15[56] 10[56] Core | AATTATCAAATATCAAAGATTAGACGCGAACT |
| 20[27] 18[9] Core | CTGAGGTCTGAGAAATCAATATATGTGAGTGCCCC |
| 11[120] 6[120] Core | TGATAATCCTCAGGAATAACAACCTCACGACG |
| 16[103] 17[103] Core | TCCAATAAATCAATAAACAATAACGGATTC |
| 0[182] 3[176] Core | CCCCAAAATCCTGTTTGATGGTCCAGCTGCATTACCCC |
| 27[56] 22[56] Core | CCAAGTACCCATATTTCGACGACAATCATAAT |
| 34[55] 31[55] Core | ATAATAACTAGCAATAGTCAGAGGAATGAAAA |
| 21[104] 23[119] Core | CGCAAGACTACCGACCAGAGGCTTTTGCAAAA |
| 12[103] 13[103] Core | AAACTAGCAAAATCTGGATTTAGTCAAAAG |
| 25[136] 28[136] Core | TATACCAGGAACGAGTTGACCTTCCGCAGACG |
| 35[9] 32[28] Core | CCCCTTATTACGCAGTATGTTAGTATC |
| 19[88] 18[88] Core | TCCTTTTGATAAGAGCATCAAGAAAACAAA |
| 33[4] 32[4] Core | CCCCTTGAGTTAAGCCCAATAGATAACCCACAAGAACCCC |
| 24[135] 29[135] Core | CCACATTCAGGCTGGCAGTAAATTGATTATAC |
| 40[71] 41[71] Core | TCATAGCCCCCTTATTGCCACCCTCAGAACCG |
| 15[120] 10[120] Core | TAGCAAAAGAGAATCGGACAGTCAAATTCGCG |
| 24[103] 25[103] Core | AAAGGAATTACGAGGAGAACGCGATTGTGA |
| 35[120] 30[120] Core | CAGTACAACGCCCACGCGTTGAAAGGCACCAA |
| 35[152] 30[152] Core | AGACAGCCCGATAGTGGAGCCTTAAACGGG |
| 41[15] 38[20] Core | CCCCACCAGAGCCACCCGTCACCACCCC |
| 18[151] 14[152] Core | CGAATTATGGCGAGTAGATTTAGATAAAAAT |
| 2[119] 0[104] Core | GGTTTTTCGCCCTTCAAGCCCGAGATAGGGTT |
| 3[88] 2[88] Core | CACCCGCCGAGCTAAAGTGAGACCCCTAAA |
| 41[72] 39[87] Core | CCACCCTCAGAGCCACTCATCGGCTAGCGTCA |
| 21[40] 24[40] Core | CTTTTTAAAAAGCCTGACAGTAGGCAACATGT |
| 11[9] 8[28] Core | CCCCCAGCAGAAGATAAAACAGATTAT |

| | |
|---|---|
| 19[152] 15[151] Core | AGCTCAACTGGGGCGCGAGCTGACAGAGCAT |
| 34[119] 31[119] Core | AGAATAGATTTTTTCACATAACCGTAAAGGCC |
| 30[198] 29[192] Core | CCCCGAGGACTAAAGACTTTTTCCCTGATAAATTGTCCCC |
| 6[87] 3[87] Core | AATCAGTGAGAATCCTATGCGCCGAACCACCA |
| 16[167] 21[167] Core | TTTTCATTATGTTTTTCCCAATAATCAAAA |
| 28[192] 27[198] Core | CCCCGTCGAAATCCGCGTCATTACCCAAATCAACGTCCCC |
| 20[71] 25[71] Core | ACAGGTCAACCCTCGTACACCGGAATAAACAA |
| 16[27] 14[9] Core | AAAAGCGTAGATATTTTAAAAGTTTGAGTAACCCC |
| 0[71] 5[71] Core | CTATTAAACGGTCACGGAGCTTGAACAGGAAC |
| 36[71] 38[56] Core | ATAGAAAAGAATCAAGAATCACCAGTAGCACC |
| 8[71] 13[71] Core | GAAATACCACAGTGCCCTTTAATGGCCGTCAA |
| 16[135] 18[120] Core | CATCAATTCTTAATTGATTACCTGAGCAAAAG |
| 36[135] 41[135] Core | CCTCATTTGTTTTAACGCTGAGACGCCAGCAT |
| 19[56] 14[56] Core | CCTTAGAATTCTGAATTATACAGTTCGTATTA |
| 23[88] 22[88] Core | AAAACCAAAATAGCGGTGTGATAAATAAGG |
| 8[39] 13[39] Core | AAATGGAGGTGAGGCGCCATTACTTTAGG |
| 29[72] 32[72] Core | GCACCCAGAACGAGCGGAGAATAAAATTAACT |
| 4[39] 9[39] Core | AACGTGCTGAGTAGACCGTTGTACATTCT |
| 36[167] 41[182] Core | CCCTCAGAGCCCGTACCTATTATGACGATTGGCCTTGATAT TCCCC |
| 1[72] 4[72] Core | GTGCCGTACCGATTTACTGCGCGTCTACAGGG |
| 3[152] 1[182] Core | CTGTCGTGGGTTCCGATCCACGCTGGTTTGCCCCAGCAGGC GCCCC |
| 8[135] 13[135] Core | GTATCGGCAGAAAAGCTCAAAAATAATCACCA |
| 9[72] 12[72] Core | AGCGTAAGCTATTAGTACGCTGAGACCTTGCT |
| 17[72] 19[87] Core | CTTTTACACATTTAACGAGTACCTTTAATTGC |
| 5[72] 8[72] Core | GGTACGCCAGGCCACCCCAGAACACGCTCATG |
| 11[56] 6[56] Core | CGCCTGCATACATTTTACCCTTCTGAGTCTGT |
| 18[87] 15[87] Core | ATTAATTATCGGGAGATAATCCTGCAGATGAT |
| 12[27] 10[9] Core | CAAATAAAATATAAAATACCGAACGAACCACCCCC |
| 11[152] 6[152] Core | TTGTATAAGCCAGTTCGGCGGAGATTAAGT |

| | |
|---|---|
| 12[71] 17[71] Core | GAACCTCATCATATTCTCGACAACAACAGTAC |
| 31[88] 30[88] Core | CAGGGAAGAGGCTTGCAAAAGAAAATCTTA |
| 1[136] 4[136] Core | GAGTTGCAGGTTTGCGTTTCCAGTCATACGAG |
| 14[151] 11[151] Core | TTTTAGAAATATTCAATGCCTGAGCAGGAAGA |
| 24[167] 29[167] Core | AGAAAGATACAAGAAGCTTGCCGAGATTTG |
| 26[119] 24[104] Core | GATGGTTTATGCGATTCAGATACATAACGCCA |
| 33[168] 36[168] Core | ATCGGTTTATGAATTTTTAGCGTAAACCGCCA |
| 37[168] 39[187] Core | CCGTACTCTTTCGGAATAAACAGTTAATGCCCCCCC |
| 24[39] 29[39] Core | AATTTAGTCATTCCTTTACGAGGAGGTTT |
| 36[39] 41[39] Core | GACAAAAGATCGATACCGGAAAACCGGAAC |
| 27[9] 24[28] Core | CCCCAATCGGCTGTCTTTCCTTAGCAG |
| 34[87] 31[87] Core | CAAAGTTAAGAAAAGTGACGGGAGCATAAAAA |
| 40[135] 38[120] Core | AGTCTCTGAATTTACCGAGCCGCCTCCTCAAG |
| 29[136] 32[136] Core | CAAGCGCGGTAATGCCCACCCTCACAATGACA |
| 38[87] 35[87] Core | GAATTAGACGTCACCGTATTTTGTGCAAAGAC |
| 5[15] 2[20] Core | CCCCATCAGAGCGGGAAGGGAAGACCCC |
| 28[71] 33[71] Core | GCGCCCAATTTACAGATCTTTCCAACCGAAGC |
| 7[152] 2[152] Core | CAACTGTTTCACAATTGTAATCATACGCGCGG |
| 12[167] 17[167] Core | AGGCTATCATCGGTTACGCAAGGTTTGACCA |
| 1[15] 1[39] Core | CCCCGCGATGGCCCACTACGTGAA |
| 28[39] 33[39] Core | GCTTATCTTGTTTATAAACAGCAAGAAA |
| 39[56] 34[56] Core | TAGCGACATTCATATGTATTCATTGAAACGCA |
| 38[187] 37[192] Core | CCCCCCTGCCTAAGGAGGTTTAGTACCCC |
| 17[40] 20[40] Core | TTTAACGTAATGGAAACTATTAATGATAGCTT |
| 26[151] 23[151] Core | AAACACCATCAGGACGTTGAGATTGTAAAATG |
| 29[168] 32[168] Core | TATCATCGATGAGGAAGAGGGTAGTTAAACAG |
| 23[9] 20[28] Core | CCCCTCTTACCAGTATAAAGCCAGACG |
| 6[151] 3[151] Core | TGGGTAACTCGAATTCCCACACAACGGGAAAC |
| 4[103] 5[103] Core | TAATGAGTGCGCTTAGAGAAGTGGCTTGCA |
| 22[151] 18[152] Core | GAGTAATTTTAGTAATGACCATATCTGCGAA |

| | |
|---|---|
| 2[176] 5[182] Core | CCCCATGAATCGGCCAGGTCATAGCTGTTTCCTGCCCC |
| 13[136] 16[136] Core | TCAATATGCCCTCATATAAAGCCTAAAGGTGG |
| 20[135] 22[120] Core | AAGACTTCGCCAGAGGTGACCTAAATTTAATG |
| 22[198] 21[192] Core | CCCCATTCATTGAATCCCCCTCATTTACCCTGACTACCCC |
| 23[152] 20[136] Core | TTTAGACTAAAAAGATTAAGAGGAAGCCCGA |
| 14[55] 11[55] Core | AATCCTTTCAACTAATACCCTCAAATTAACAC |
| 0[135] 5[135] Core | TATAAATCCTGCCCGCTATTGGGCTCCCCGGG |
| 14[119] 11[119] Core | CCTGAGTAAGGCCGGAATGAACGGACCCCGGT |
| 28[103] 29[103] Core | TGAAAGAGTTTTCATTTATCCTGTACACTA |
| 32[135] 37[135] Core | ACAACCATACTACAACCAGTTTCAAGAGGGTT |
| 33[72] 36[72] Core | CCTTTTTACCAGAAGGAAAGAAACCACAATCA |
| 8[192] 7[198] Core | CCCCGTAGATGGGCGCATGCGGGCCTCTTCGCTATTCCCC |
| 13[4] 12[4] Core | CCCCATTGAGGAAGGTTATCTCAACAGTTGAAAGGACCCC |
| 19[120] 14[120] Core | GCTTAGAGCTACTAATTACCAAGTATGCAATG |
| 21[168] 24[168] Core | ATCAGGTCAATGCTTTGTCCAATATTACAGGT |
| 7[9] 4[15] Core | CCCCGTAATAACATCACTTGCCTTTCCTCGTTAGACCCC |
| 27[88] 26[88] Core | CGTTTTTAGACAGATCTTTAATCCCTGTTT |
| 29[40] 32[40] Core | TGAAGCCTTTACAAAAACGTCAAAGTAATTGA |
| 20[103] 21[103] Core | TCAAAGCGAACCAGATGATGCAAATCCAAT |
| 25[168] 28[168] Core | TACGTTAATGAATAAGCCGGATATACCTGCTC |
| 34[198] 33[192] Core | CCCCTCTTTCCAGACGTTAGTAAATCAGCTTGCTTTCCCC |
| 32[192] 31[198] Core | CCCCCGAGGTGAATTTCCAACGGCTACAGAGGCTTTCCCC |
| 3[120] 1[135] Core | TGCGCTCAAAAAGAATCCGCCTGGCCCTGAGA |
| 21[136] 24[136] Core | AATATATTTCATCTTCGGGTAATATAGGAATA |
| 26[87] 24[72] Core | ATCAACAAGCTAATGCCATAGTAAGAGCAACA |
| 35[88] 34[88] Core | ACCACGGAACCGTAAACTAAAGGAGCCGAA |
| 30[151] 27[151] Core | TAAAATACAAACAAAGGAACGAGGATCAAGAG |
| 17[136] 19[151] Core | CGCGCAGATCATTTCACTGAATATAATGCTGT |
| 37[136] 40[136] Core | GATATAAGATTAAGAGGGGGTCAGGAAAGCGC |
| 39[88] 38[88] Core | GACTGTAGTTTTGATGCGGGGTTCATTTGG |

| | |
|---|---|
| 26[198] 25[192] Core | CCCCAACAAAGCTGCTCATTCAGTAAAACGAACTAACCCC |
| 33[40] 36[40] Core | CAATGAAAGGAATACCAGAAAATAGCGCCAAA |
| 37[72] 40[72] Core | ATTATCACGCCAGCAATTTGCCTTATTTTCGG |
| 31[56] 26[56] Core | TAGCAGCCTAGCAAGCGATTAGTTAAGAAAAA |
| 32[39] 37[39] Core | GCGCTAACAAACGTCAAAAGAAGGGAAGGT |
| 21[4] 20[4] Core | CCCCATTTATCAAAATCATAGAAGAGTCAATAGTGACCCC |
| 32[103] 33[103] Core | GGTCGCTGCGCATTAAAGCAGATAATTGCG |
| 16[71] 18[56] Core | GATTATACGGATTAGAAATTTCATTTGAATTA |
| 10[87] 7[87] Core | TTGAATGGAATACGTGAGGAAAAAATATTACC |
| 10[198] 9[192] Core | CCCCTAAAATTCGCATTAAATTTAGGTCACGTTGGTCCCC |
| 38[55] 35[55] Core | ATTACCATACGGAAATGTTTACCACATACATA |
| 15[9] 12[28] Core | CCCCCATTATCATTTTGCGGAACTTGG |
| 22[55] 19[55] Core | TACTAGAACCTCCGGCAACATAGCTAATTTTC |
| 30[87] 27[87] Core | CCAACGCTCTACAATTCGTAGGAAAAGCAAGC |
| 29[104] 32[104] Core | AAACACTCGAAAGAGGCAGGGAGTATATATTC |
| 22[87] 20[72] Core | CGTTAAATGTAAATGCCCGGAAGCAAACTCCA |
| 36[103] 37[103] Core | ACCCATGTATAAGTTACTTGAGCTTGCTCA |
| 7[120] 2[120] Core | CAGGCAAAATAAAGTGCTAGAGGAGCCAGGGT |
| 32[71] 37[71] Core | GAACACCCAACATATAAAACCGAGAAAGGTGA |
| 7[56] 2[56] Core | GCCTTGCTATGGTTGCGATTTTAGCGGGGAAA |
| 0[103] 1[103] Core | GAGTGTTGTTCCAGTAATCGGAAGGGCAAC |
| 14[87] 11[87] Core | AGACTTTAACATTTGAAAAGCATCAGCCAGCA |
| 8[27] 6[9] Core | TTACTAAAAGGGAGCAATACTTCTTTGATTACCCC |
| 9[136] 12[136] Core | TCTCCGTGGAACGCCACCCAAAAAAGTCTGGA |
| 41[104] 39[119] Core | ACCAGAACCACCACCAGTTCCAGTAGTGTACT |
| 8[103] 9[103] Core | GCTTTCCGTTGCAACGCACAGACCATCAAC |
| 39[152] 34[152] Core | GTAACAGTACCGCCAGGAATAGGGGATTTT |
| 29[4] 28[4] Core | CCCCTTTAGCGAACCTCCCGTAAGAACGCGAGGCGTCCCC |
| 33[104] 36[104] Core | AATAATAAAAGGAACACACTGAGTCAATAGGA |
| 2[55] 0[40] Core | GCCGGCGACAAATCAAACTCCAACGTCAAAGG |

| | | |
|---|---|---|
| 31[120] 26[120] Core | GCTTTTGCTAAGGGAACCCCCAGCGGGCTTGA | |
| 0[39] 5[39] Core | GCGAAAAGGAGCGGAGAAAGGAGCTAAACA | |
| 40[103] 41[103] Core | TACATGGCCGCGTTTCACCCTCAGAGCCGCC | |
| 1[40] 3[55] D2 | CCATCACCACGTGGCGGCGCTAGGGCGCTGGC**TTTTAGGTAAATTTTGATTGTGAGGAAG** | **D2** |
| 13[104] 15[119] D2 | GGTGAGAAATGTGTAGAGGCAAGGCAAAGAAT**TTTTAGGTAAATTTTGATTGTGAGGAAG** | **D2** |
| 13[40] 15[55] D2 | AGCACTAAGCCCGAACCCACCAGAAGGAGCGG**TTTTAGGTAAATTTTGATTGTGAGGAAG** | **D2** |
| 1[104] 3[119] D2 | AGCTGATTTTTTCACCCTCACATTAATTGCGT**TTTTAGGTAAATTTTGATTGTGAGGAAG** | **D2** |
| 25[104] 27[119] D2 | ATTACCTTAATTTCAAGAACGGTGTACAGACC**TTTTAGGTAAATTTTGATTGTGAGGAAG** | **D2** |
| 25[40] 27[55] D2 | AGTAATTCCATCCTAAAAGAACGGGTATTAAA**TTTTAGGTAAATTTTGATTGTGAGGAAG** | **D2** |

### Supplementary Figures

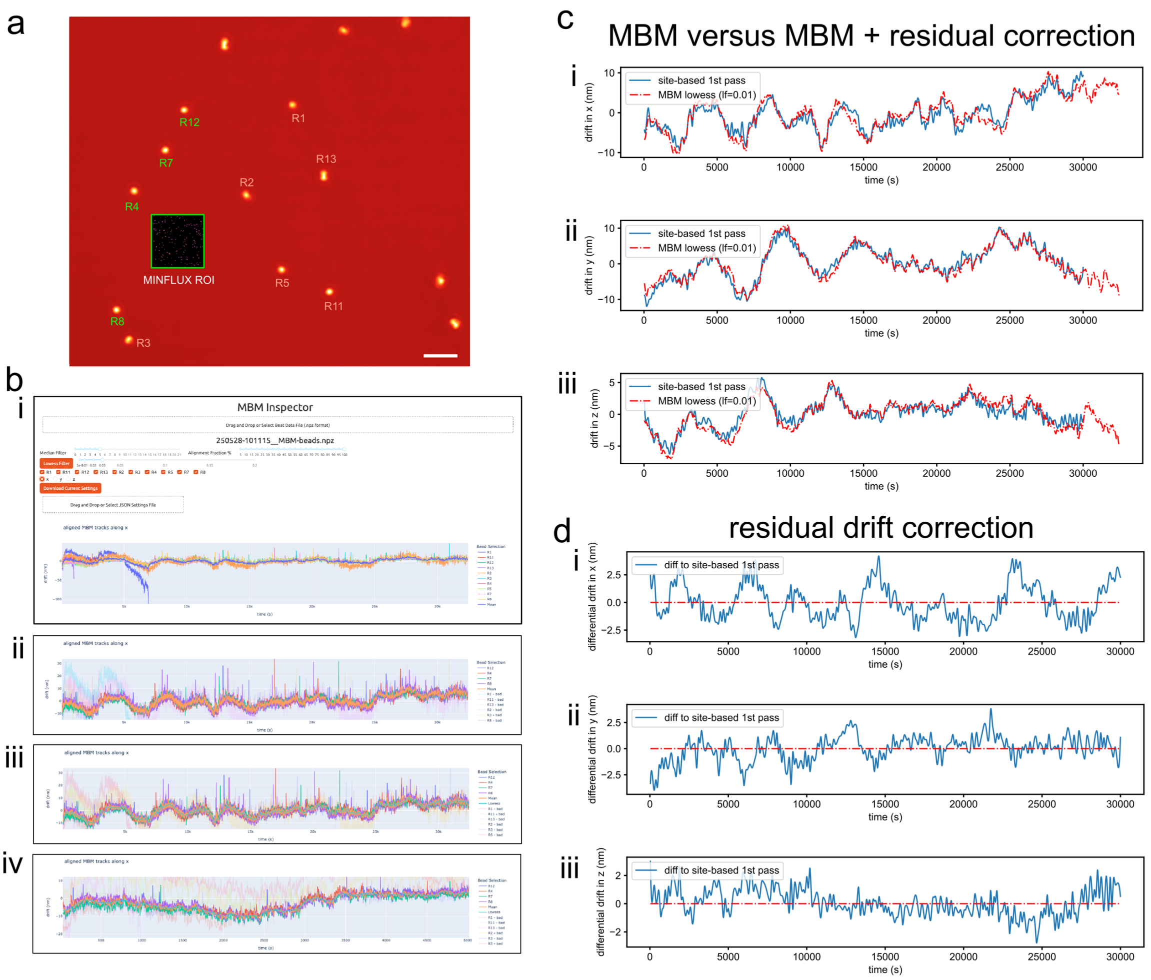

Supplementary Figure 1. Processing of MBM data. **a**. A confocal reflection scan of a sample containing DNA origami structure O1 (MINFLUX ROI shown as overlay) and surrounding gold nanoparticles selected for MBM tracking. MBM beads are labeled R<number>, in the example shown bead R13 exhibits a structure compatible with two adjacent particles being present. Beads selected for calculating the mean MBM trajectory are labeled green, those deselected are labeled in red. **b**. **i**. Screenshot of the MBM Inspector app after loading MBM bead trajectories and displaying the trajectories in the x-coordinate. Note the trajectory of bead R1 noticeably diverging. **ii**. Trajectories in the x direction after deselecting diverging beads, deselected trajectories appear faint and beads are marked "bad" in the legend. **iii**. Corresponding selected trajectories in y and **iv**. z-directions. Note the overlayed mean trajectory (calculated only from selected beads) and in iii and iv also shown are overlaid lowess smoothed mean trajectories. **c**. Resulting mean MBM trajectories (blue lines) compared to total estimated drift (red, representing mean MBM trajectory + residual drift from DNA origami based correlated site-dispersion estimate) in x (**i**), y (**ii**) and z (**iii**) directions. **d**. Corresponding residual drift trajectories, i.e. the difference curves between blue and red lines in c, shown in x (**i**), y (**ii**) and z (**iii**) directions. Scale bar a: 2 µm.

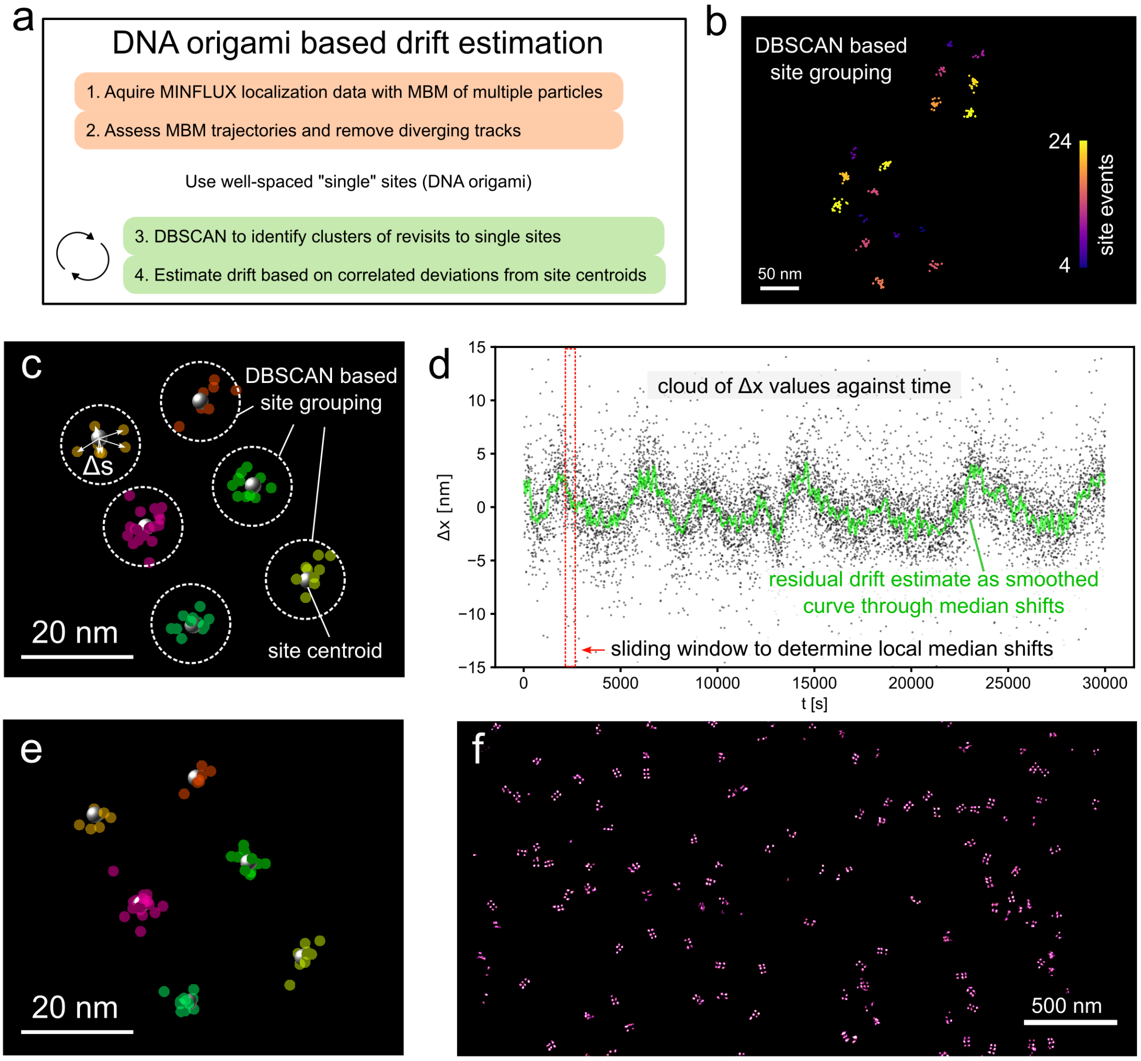

Supplementary Figure 2. Residual drift estimation by identifying correlated site-dispersion. **a**. Schematic of the data processing pipeline. The green panels designate the main steps of the residual drift estimation approach. **b**. Groups of localizations originating from the same anchor site were identified by DBSCAN clustering with an epsilon of 10-15 nm. The result of DBSCAN grouping is shown here with site groups from O2 DNA origami structures colored by the number of localizations (here containing between 4 and 24 localizations). **c**. For each site group the centroid (grey spheres) was determined from all localizations in the group and the dispersion of localizations around the site centroid was determined by calculating the shift Δs from the centroid position for each localization. Color indicates localizations in the same site group. **d**. The shifts in each coordinate direction (Δx, Δy, Δz) plotted against acquisition time exhibit correlated shifts (correlated site-dispersion) indicated by regions of high point density as illustrated here for Δx. To estimate the residual drift value indicated by these time shifts the median shift in a sliding time window is determined and the sequence of drift values obtained in this way is finally smoothed to obtain a residual drift trajectory (green line). Typical window sizes are between 50 to 400 s to ensure enough localizations fall into a time window for a robust estimate. **e**. After determining residual drift trajectories in x, y and z directions these are subtracted from the input coordinates (panel c) to obtain corrected coordinates. The corrected coordinates are visibly tighter packed around the site centroids. **f**. Overview rendering of the dataset from which data in b-e was extracted after removal of residual drift.

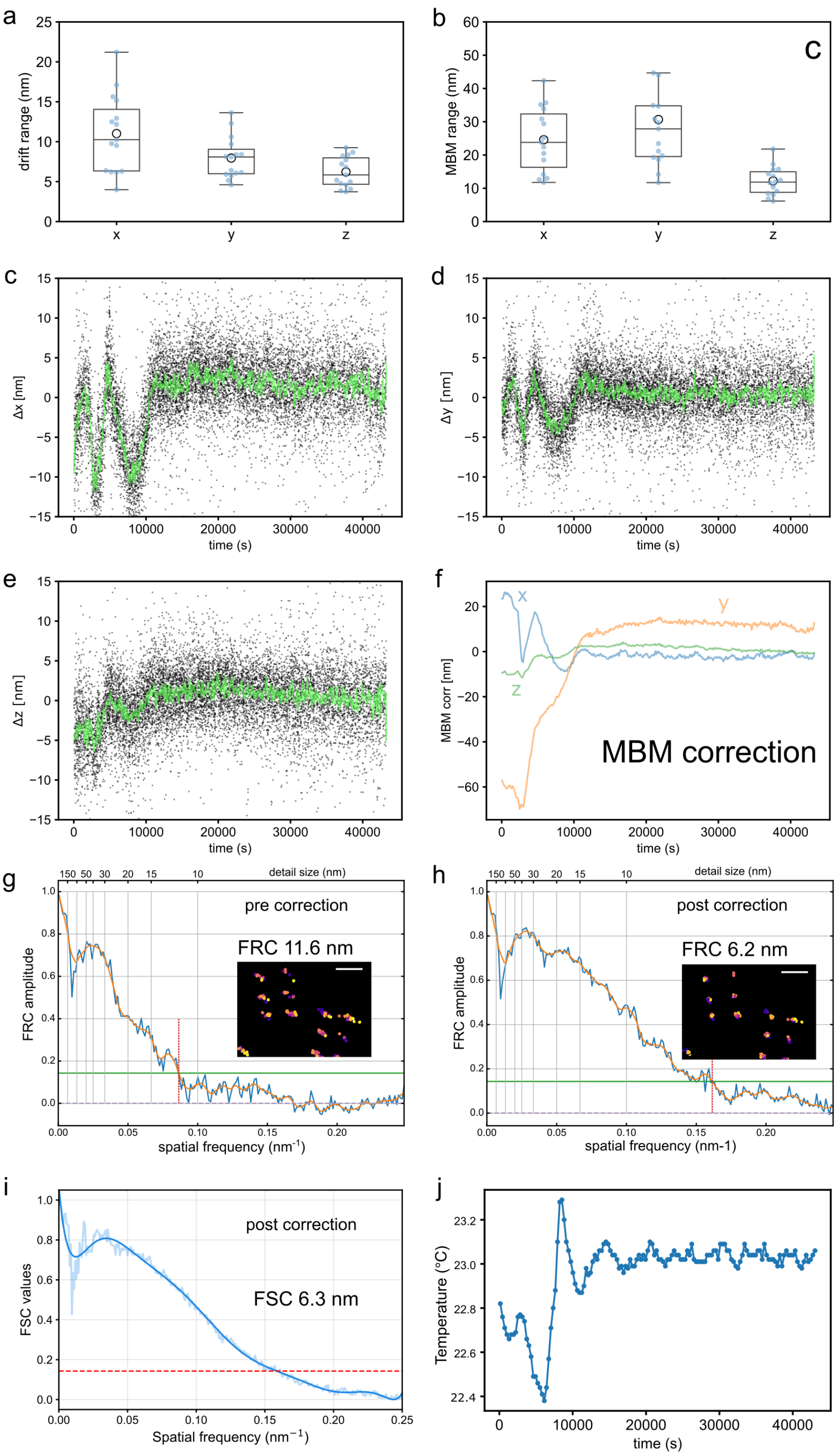

Supplementary Figure 3. Drift magnitude statistics and a sample dataset with more pronounced residual drift. **a**. The residual drift range magnitude quantified as the difference between the maximal and minimal residual drift over an acquisition duration of 30K s was between ~10 nm (median in x) and ~6 nm (median in z), (n=15 data sets from N=6 independent replicates). **b**. Associated median MBM recorded drift ranges were between ~28 and ~12 nm (n=15 data sets from N=6 independent replicates). **c, d, e**. Residual drift trajectories (in x, y, and z directions, respectively) from an example acquisition with a more marked residual drift over the first ~12000 s. **f**. Associated MBM trajectories that exhibit substantial amplitudes (~80 nm in the y direction). **g**. Before correcting residual drift the Fourier ring correlation (FRC), a measure of lateral resolution, was 11.6 nm with MBM correction only. **h**. FRC resolution improved to 6.2 nm once the residual drift was additionally corrected. For FRC calculations shown in g and h, data from the initial 12000 s were used for rendering. **i**. The Fourier Shell Correlation (FSC), a measure of 3D resolution, after residual correction, was 6.3 nm for a ROI from this dataset. **j**. Temperature changes at the microscope stand during the acquisition of the data series shown in c-f. Note the initial temperature swing by ~0.8 °C. The more stable temperature from ~15000 s onwards was associated with reduced MBM drift and similarly low residual drift. Scale bars 50 nm.

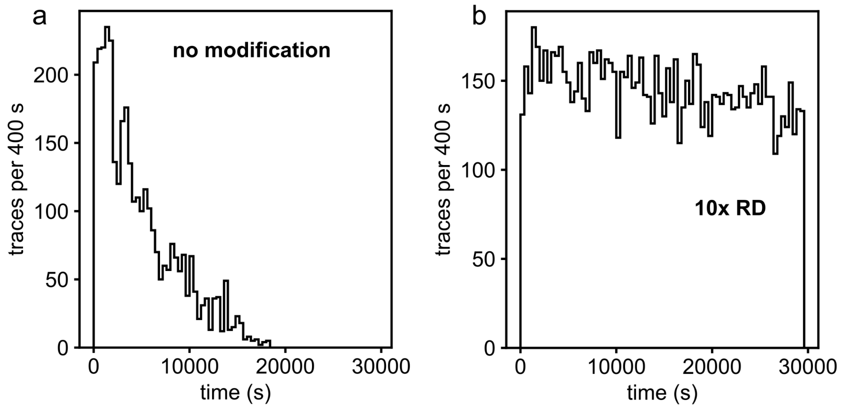

Supplementary Figure 4. Exemplary localization rate time courses during 3D MINFLUX acquisition with DNA origami structure O2. **a**. Localization rate quantified as number of complete and valid MINFLUX traces per 400 s from an acquisition where unmodified anchor strands containing a single P1* domain were probed with P1s imagers (see also Table 1 in main text). Note the relatively rapid decay of localization rate so that the acquisition had to be halted at ~18000 s. This reduction in localization rate represents extensive site-loss. **b**. By contrast, when using the 10xRD P1 docking strands (Table 1) attached to anchor strands on O2 and then equally probed with P1s imagers, localization rate remains almost undiminished through 30000 s.

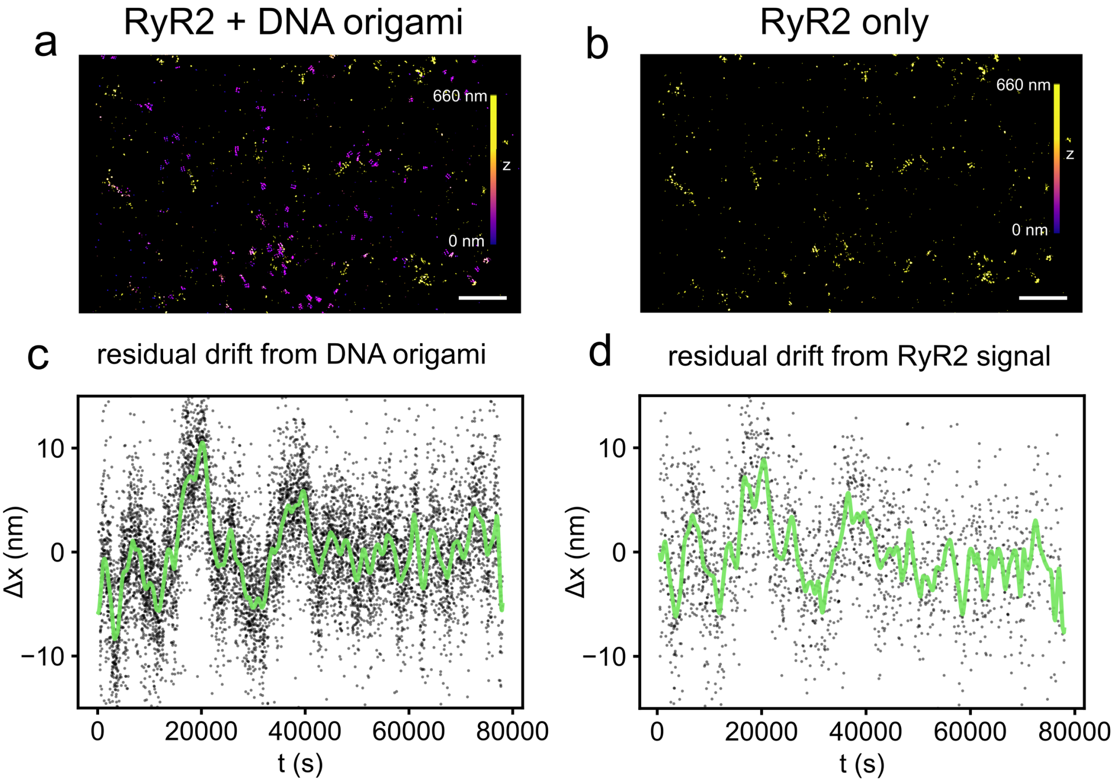

Supplementary Figure 5. Residual drift correction estimates from DNA origami versus estimates obtained directly from protein target signals with repeat domain sdABs. **a**. Overview 3D MINFLUX image of a cardiac tissue section containing DNA origami structure O1 and sdABs against tagFP targeting PAtagRFP residues on RyR2 proteins. The DNA origami structures reside almost exclusively at low z levels < 300 nm (blue/magenta) whereas RyR2 signals were at z levels > 350 nm (yellow). **b**. Overview after filtering z coordinates to only include RyR2 signal. **c**. Correlated shifts and estimated residual drift trajectory in the x direction only using the DNA origami sites. **d**. Equivalent shifts and trajectory estimated only from sites on RyR2. In this sample the number of RyR2 sites is smaller than the number of DNA origami sites. Nevertheless, the trajectory estimates are comparable. Scale bars 1 µm.
